# General dimensions of human brain morphometry inferred from genome-wide association data

**DOI:** 10.1101/2021.10.22.465437

**Authors:** Anna E. Fürtjes, Ryan Arathimos, Jonathan R. I. Coleman, James H. Cole, Simon R. Cox, Ian J. Deary, Javier de la Fuente, James W. Madole, Elliot M. Tucker-Drob, Stuart J. Ritchie

## Abstract

**Background:** Understanding the neurodegenerative mechanisms underlying cognitive declines in the general population may facilitate early detection of adverse health outcomes in late life. This study investigates biological pathways shared between brain morphometry, ageing, and cognitive ability.

**Methods:** We develop Genomic Principal Components Analysis *(genomic PCA)* to model general dimensions of variance in brain morphometry within brain networks at the level of their underlying genetic architecture. With genomic PCA we extract genetic principal components (PCs) that index global dimensions of genetic variance across phenotypes (unlike ancestral PCs that index genetic similarity between participants). Genomic PCA is applied to genome-wide association data for 83 brain regions which we calculated in 36,778 participants of the UK Biobank cohort. Using linkage disequilibrium score regression, we estimate genetic overlap between brain networks and indices of cognitive ability and brain ageing.

**Results:** A genomic principal component (PC) representing brain-wide dimensions of shared genetic architecture accounted for 40% of the genetic variance across 83 individual brain regions. Genomic PCs corresponding to canonical brain networks accounted for 47-65% of the genetic variance in the corresponding brain regions. These genomic PCs were negatively associated with brain age (*r_g_* = −0.34). Loadings of individual brain regions on the whole-brain genomic PC corresponded to sensitivity of a corresponding region to age (*r* = - 0.27). We identified positive genetic associations between genomic PCs of brain morphometry and general cognitive ability (*r_g_*= 0.17-0.21).

**Conclusion:** These results demonstrate substantial shared genetic etiology between connectome-wide dimensions of brain morphometry, ageing, and cognitive ability, which will help guide investigations into risk factors and potential interventions of ageing-related cognitive decline.

## 1. Introduction

Progressive ageing-related neurodegenerative processes occurring across micro to macro scales of the human brain are well-documented within otherwise healthy adults, and are linked to ageing-related declines in multiple domains of cognitive function [1–3]. Understanding the biological processes underlying these links is paramount for identifying mechanisms of cognitive ageing that can ultimately be targeted by intervention. The human brain is a complex network of interconnected regions (the ‘connectome’) [4, 5], components of which are interrelated with one another [6, 7], age unevenly over time [8], and may be differentially relevant to adult cognitive ageing [1–3]. Whereas considerable attention has been devoted separately to the genetic architecture of human brain morphometry [9–11] and the genetic architecture of adult cognitive ability [12], relatively less work has been devoted to scaffolding investigations of the genetic architecture of human brain morphometry onto the well-established network organization of the brain (although see [13] for a recent exception), or to investigating how genetic links between components of human brain networks relate to ageing and cognition. Such investigations have the potential to provide insights into the etiology of neurocognitive ageing.

Specifically, our genome-wide study builds on a previous study that investigated dimensions of brain morphometric variation (i.e., principal components underlying brain volumetric measures) across the human connectome in a large scale cohort (*N* = 8,185) [3]. This phenotypic study implicated well-studied macroscopic brain networks in cognitive ageing, whereby connectome aging varied alongside those dimensions of morphometric variation. Brain volumes in the central executive network tended to be most sensitive to age (i.e., cross-sectionally correlated with age) and, albeit its small size, the central executive was highlighted to play a disproportionate role in late-life cognitive ability.

Here, we hypothesise that morphometric network organisation, as described by Madole et al. [3], corresponds to morphometric variation captured by genome-wide data (pre-registration: https://osf.io/7n4qj). We suggest that a dissimilar organisation of phenotypic and genetic brain architecture would contradict the neurobiological validity of canonical brain networks (a similar organisation would be consistent with a measurable genetic foundation of brain networks). This hypothesis relies on evidence that patterns of brain morphometry and its organisation are highly heritable [10, 11, 14, 15]. Cheverud originally speculated that “If genetically and environmentally based phenotypic variations are produced by similar disruptions of developmental pathways, genetic and environmental correlations should be similar.” [16]. Strong correspondence between phenotypic and genetic correlations was recently demonstrated for a wide range of morphometric human traits (for example, height and body mass index) in the UK Biobank cohort [17], and we therefore expect that phenotypic brain network structures should mirror the structure of genetic correlations within the same networks.

We consider the same ‘canonical’ brain networks as Madole et al. [3], using common, but not indisputable, definitions of the exact regions comprising them [5, 18, 19]. These brain networks have been characterised embracing a whole-brain perspective, considering existing literature describing synchronised (i.e., correlated) regional activity in functional MRI data [3], in addition to converging evidence from other modalities (i.e., structural MRI and lesion-based mapping [7, 20, 21]). Among the most reported networks are the central executive, default mode, salience, and multiple demand networks. Specific network characterisations considered in this study are displayed in Fig. 1 and listed in STable 2.

**Fig. 1.**
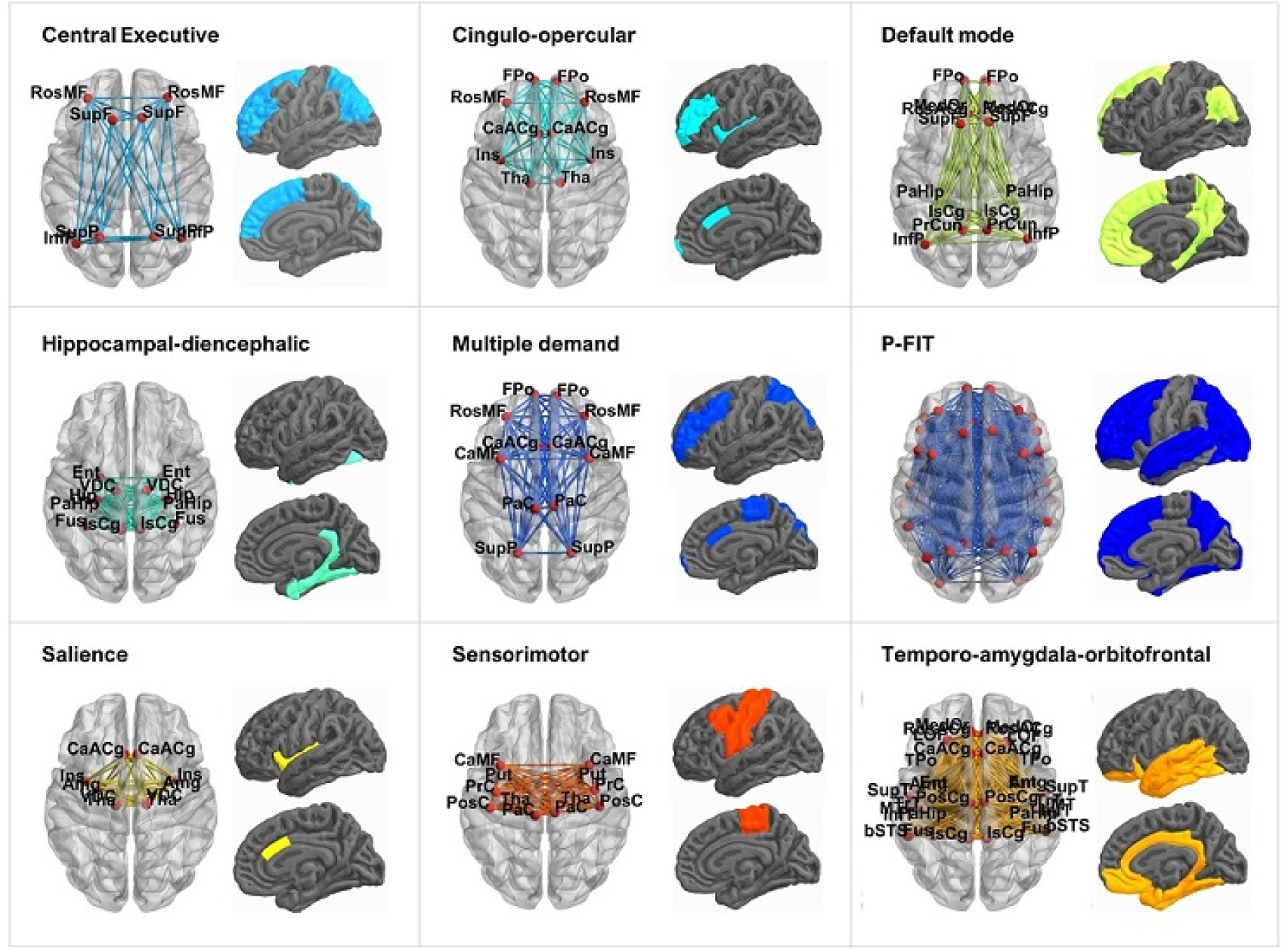
Nine canonical brain subnetworks. The networks were visualized with the BrainNet Viewer (http://www.nitrc.org/projects/bnv/) [61]. Regions of interest were visualised using scripts by Dr. Colin Buchanan (University of Edinburgh). Included brain regions and their abbreviations are listed in STable 2.

Brain networks are theorised to integrate information across the brain and, collectively, to give rise to cognitive functions. The central executive network is thought to underpin higher-level cognitive functions, including attention and working memory processes [21, 22]; whereas the default mode network is associated with internally directed and abstract thought [23]. The salience network is thought to detect salient sensory cues [24], helping to integrate executive and default functions [22, 25]. Mental processes that organise multiple cognitive requirements into a series of successive cognitive tasks are thought to be associated with the multiple demand network [26].

Here, we use genome-wide association data summarising genetic correlates of grey matter volumes to model morphometric network structures. We focus on brain volumes because they are highly heritable [14], and are measured independent of mental processes during MRI scanning (compared with functional MRI). Grey matter volume was demonstrated to be a strong and robust predictor of general cognitive ability [27, 28], it reflects atrophy; an important indicator of ageing and health outcomes [29], and, as discussed above, regional brain volumes have been shown to capture dimensions of morphometric variation implicated in aging and cognitive ability [3].

In order to model macroscopic brain networks using genome-wide association data, in this pre-registered study (https://osf.io/7n4qj), we present our novel statistical genetics method ‘genomic PCA’ (genomic Principal Component Analysis). With genomic PCA we extract genetic principal components (PCs) that index global dimensions of genetic variance across phenotypes (unlike ancestral PCs that index genetic similarity between participants), the human structural connectome. We mirror previous phenotypic analyses to ensure comparability with results in Madole et al. [3], that is, we estimate genetic associations between genetic PCs across brain network structures and both cognitive ability and ‘brain age’ [30]. Characterising genetic links between general dimensions of brain organisation, aging and cognitive ability will help guide investigations into risk factors, biological mechanisms, and potential interventions of ageing-related cognitive decline.

## 2. Methods

The considered UK Biobank sample consisted of 36,778 participants (54% females) with available neuroimaging data and had an average age of 63.3 years at neuroimaging visit (range from 40.0 to 81.8 years). Our methodological approach followed four major analysis steps as displayed in Fig. 2. First, we calculated 83 genome-wide association study (GWAS) summary statistics for 83 regional volumes that served as input data. Polygenic effects were fitted in a linear mixed model using REGENIE [31]. Second, we calculated genetic correlation matrices indicating genetic overlap between regional brain volumes using the GenomicSEM software [32]. Genetic correlations formed the basis for subsequent analyses; they are contrasted with phenotypic correlations in Section 3.1. Third, we extracted the first principal component (PC) from genetic correlation matrices using principal components analysis (PCA). The genetic PCs were calculated for the whole brain, as well as canonical brain networks; they represent general dimensions of shared genetic morphometry between regions included in either the whole brain, or the brain networks (Section 3.2). Based on Pearson’s correlations and Tucker congruence coefficient, we compared genetic correlation structures with phenotypic correlation structures between the 83 regional volumes (Section 3.3). We also tested whether the relative ordering of phenotypic and genetic PC loadings correlated with indices of a regions sensitivity to age (i.e. cross-sectional volume-age correlations; Section 3.4). Finally, we presented a novel method to summarise shared morphometric variance within brain networks on a genome-wide level in sets of univariate summary statistics (i.e., genetic PCs underlying multiple brain volumes; see Supplementary Methods). These univariate summary statistics can be viewed as a summary-based method of computing GWAS summary statistics that would be obtained from a GWAS on individuals’ scores on the underlying genetic PCs. We used those summary statistics to represent general dimensions of brain organisation across the whole brain and nine canonical networks at the level of their underlying genetic architecture. We then quantified genetic correlations between brain network structures and both general cognitive ability [32] (Section 3.5) and brain age (Section 3.6). Detailed descriptions of the UK Biobank data used in this study and the study design can be found in the Supplementary Methods. Our analysis code is displayed at https://annafurtjes.github.io/Genetic_networks_project/.

**Fig. 2.**
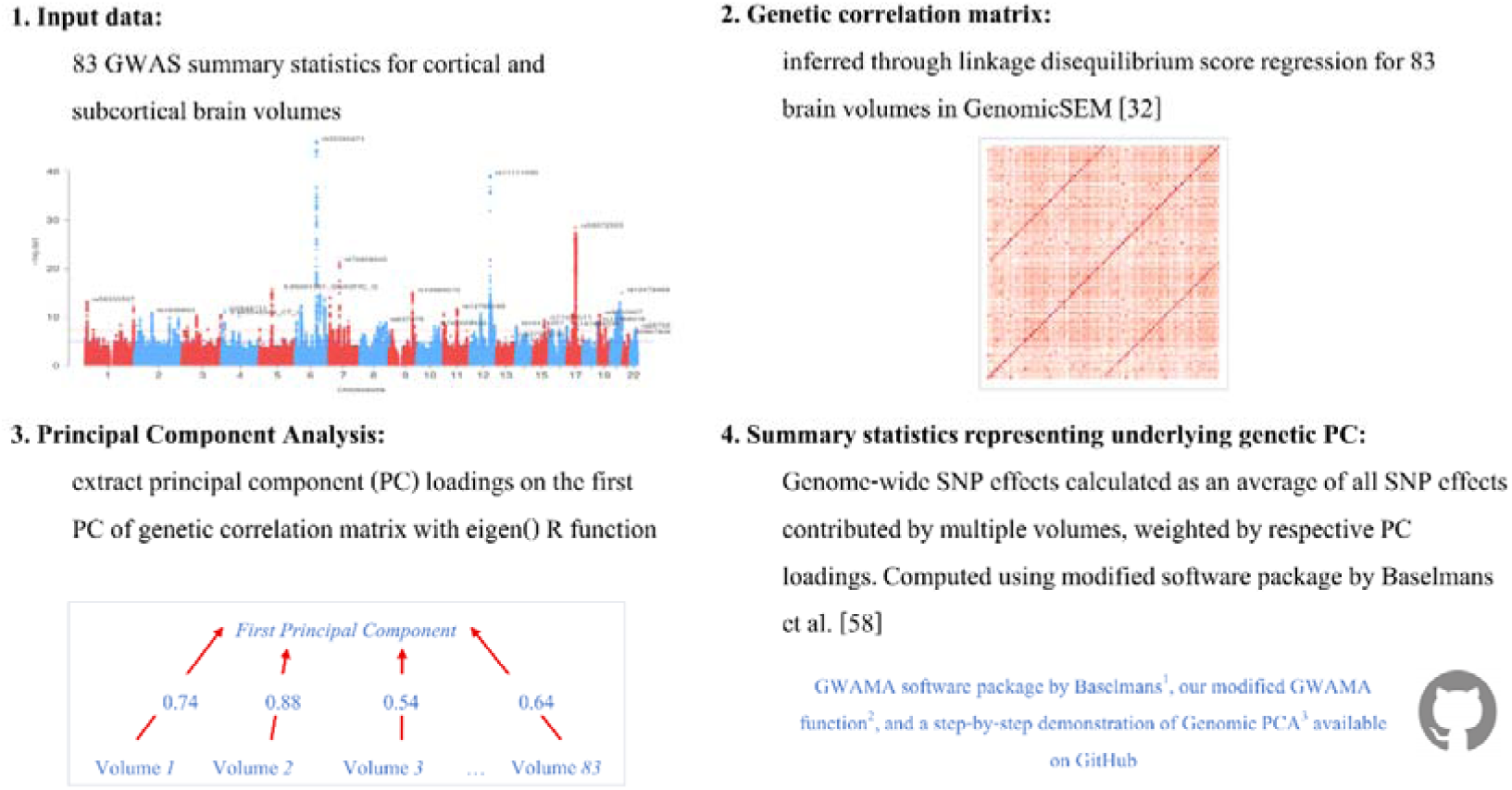
Four-step procedure to obtain statistical representation of genetic brain network structure from GWAS summary statistics. (1) GWAS summary statistics for 83 grey-matter volumes in UK Biobank from European ancestry were used as input data (*N* = 36,778). They were calculated as described in Methods and are publicly available. (2) Linkage disequilibrium score regression (LDSC) was used to infer genetic correlations between 83 brain volumes using GenomicSEM [32]. (3) Genetic correlations are analysed using PCA to derive PC loadings on the first PC, representing an underlying dimension of shared morphometry. (4) We developed a method to derive univariate summary statistics for a genetic PC of multiple GWAS phenotypes (derived from samples of unknown degrees of overlap). A genetic PC underlying several brain volumes is interpreted throughout the manuscript to index general dimensions of regionally shared morphometry. Genome-wide SNP effects are calculated as an average of all SNP effects contributed by multiple phenotypes, weighted by their respective PC loadings. Standard errors are computed using a method that corrects for sample overlap, as estimated by LDSC. We have validated this novel approach in an independent set of GWAS summary statistics [59]. All software we used is available on https://github.com/. ^1^ The software by Baselmans, Jansen [58], containing the GWAMA function is available at https://github.com/baselmans/multivariate_GWAMA/. ^2^ Our modified version of the GWAMA function is at https://github.com/AnnaFurtjes/Genetic_networks_project/blob/main/my_GWAMA_26032020.R and ^3^ a step-by-step demonstration of genomic PCA is at https://annafurtjes.github.io/genomicPCA/.

## 3. Results

### 3.1 Genetic correlations between brain-wide volumes recapitulated phenotypic correlations

On a phenotypic level of analysis, correlations between the 83 brain volumes were obtained using Pearson’s correlations from volumetric phenotypes that were residualised for age (mean = 63.3, range = 40.0-81.8 years) and sex (54% females). On a genetic level of analysis, we calculated GWAS summary statistics to get genome-wide associations (*N* = 36,778) for 83 cortical and subcortical grey-matter volumes (Fig. 2.1). SNP-heritability estimates ranged between 7% (*SE* = 0.07) for the frontal poles and 42% (*SE* = 0.04) for the brain stem (mean = 0.23, *SD* = 0.07; Fig. 3A).

**Fig. 3.**
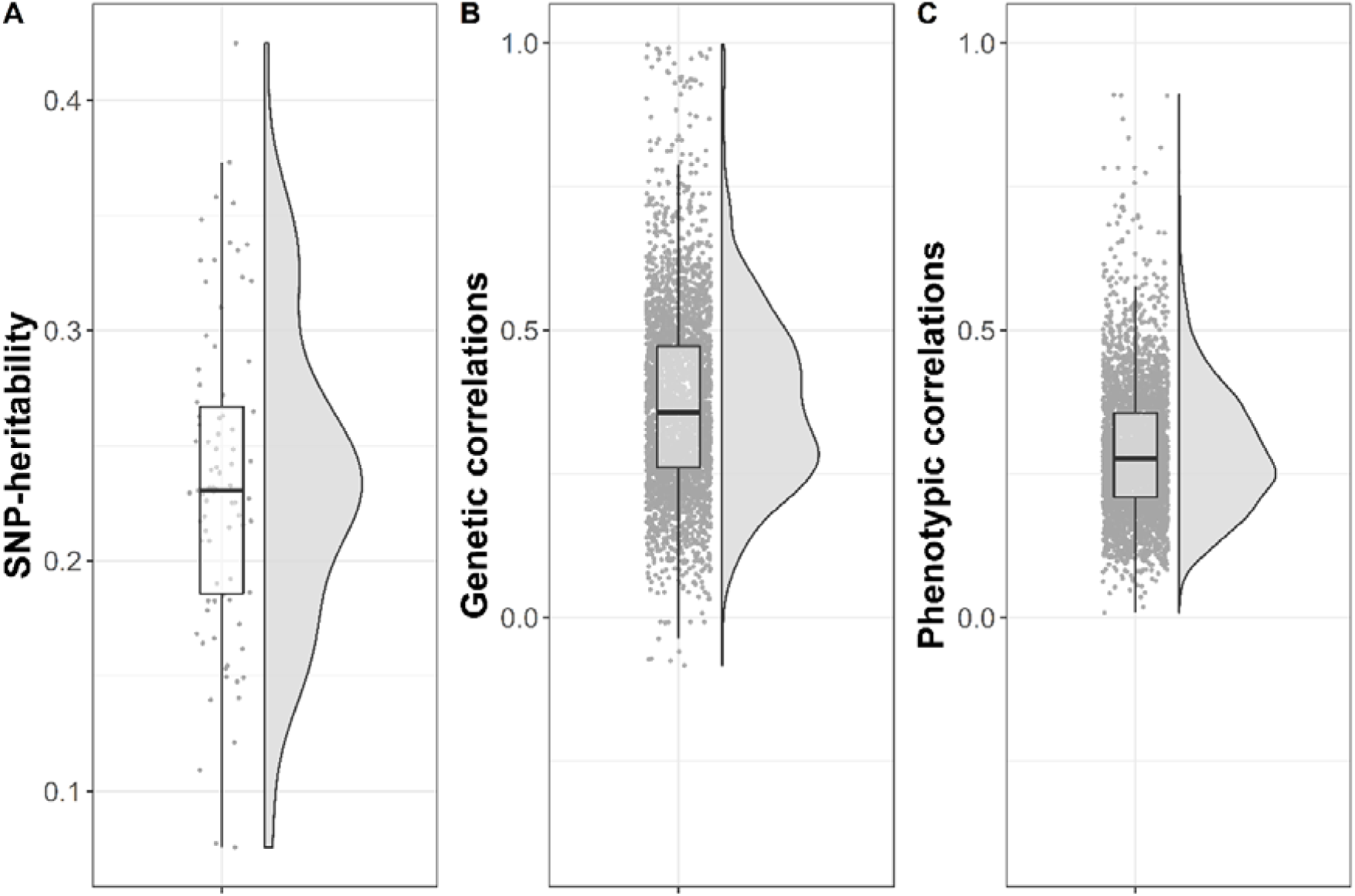
**(A)** Distribution of SNP-heritability estimates for 83 regional grey-matter volumes inferred through univariate LDSC. **(B)** Distribution of genetic correlations among 83 regional grey-matter volumes inferred through between-region LDSC. This figure only depicts between-region correlations but not the very high genetic inter-region correlations between regions and their homologous counterpart in the opposite hemisphere (excluding brain stem). **(C)** Distribution of phenotypic correlations among 83 regional grey-matter volumes inferred through Pearson’s correlations. The raincloud plots were created based on code adapted from Allen et al. [62].

The GWAS summary statistics enabled the calculation of genetic correlations between 83 volumes through linkage disequilibrium score regression (LDSC) [33]( 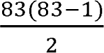 = 3403 between-region correlations; Fig. 2.2). All bilateral regions were almost perfectly correlated with the corresponding contralateral region. Between-region genetic correlations ranged from *r_g_* = −0.08 (*SE* = 0.09) between right frontal pole and left pallidum, to *r_g_* = 0.87 (*SE* = 0.08) between left middle temporal and left inferior temporal (Fig. 3B, SFig. 1). Corresponding standard errors ranged between 0.01 and 0.03 (mean = 0.014; *SD* = 0.002). Genetic correlations within canonical networks are provided in SFig.s 2-10.

A positive and large association (*r* = .84; *b* = 0.60; SE = 0.007, *p* < 2 x10^-16^, *R^2^* = 70%) was obtained between 3403 phenotypic and 3403 genetic correlations (Fig. 3&5A), indicating that the same regions, that had strongly correlated phenotypic volumes, were also genetically correlated. Phenotypic correlations were exclusively positive, as were 3,392 of 3,403 genetic correlations; the 11 (0.32%) negative genetic correlations were close to zero (smallest *r_g_* −0.083; Fig.3).

### 3.2 PCs of shared genetic variance across the whole-brain and canonical networks

Distributions of phenotypic PC loadings are in Fig. 4A (descriptive statistics for phenotypic shared morphometry in STable 3). On a genetic level of analysis, we extracted PCs from genetic correlation matrices. The first genetic whole-brain PC explained 40% of the genetic variance across 83 regional volumes - slightly larger than the 31% explained by the first phenotypic whole-brain PC. The second genetic whole-brain PC accounted for 6.7% of the total genetic variance; that is, 17% of the variance explained by the 1^st^ genetic PC (SFig. 20). We obtained loadings on this first genetic PC for each regional volume, quantifying how well an individual volume mapped onto the underlying dimension of shared morphometry across the whole brain. Their distribution ranged between 0.30 and 0.81 (mean = 0.62, *SD* = 0.13, median = 0.65; Fig. 4, STable 3a).

**Fig. 4.**
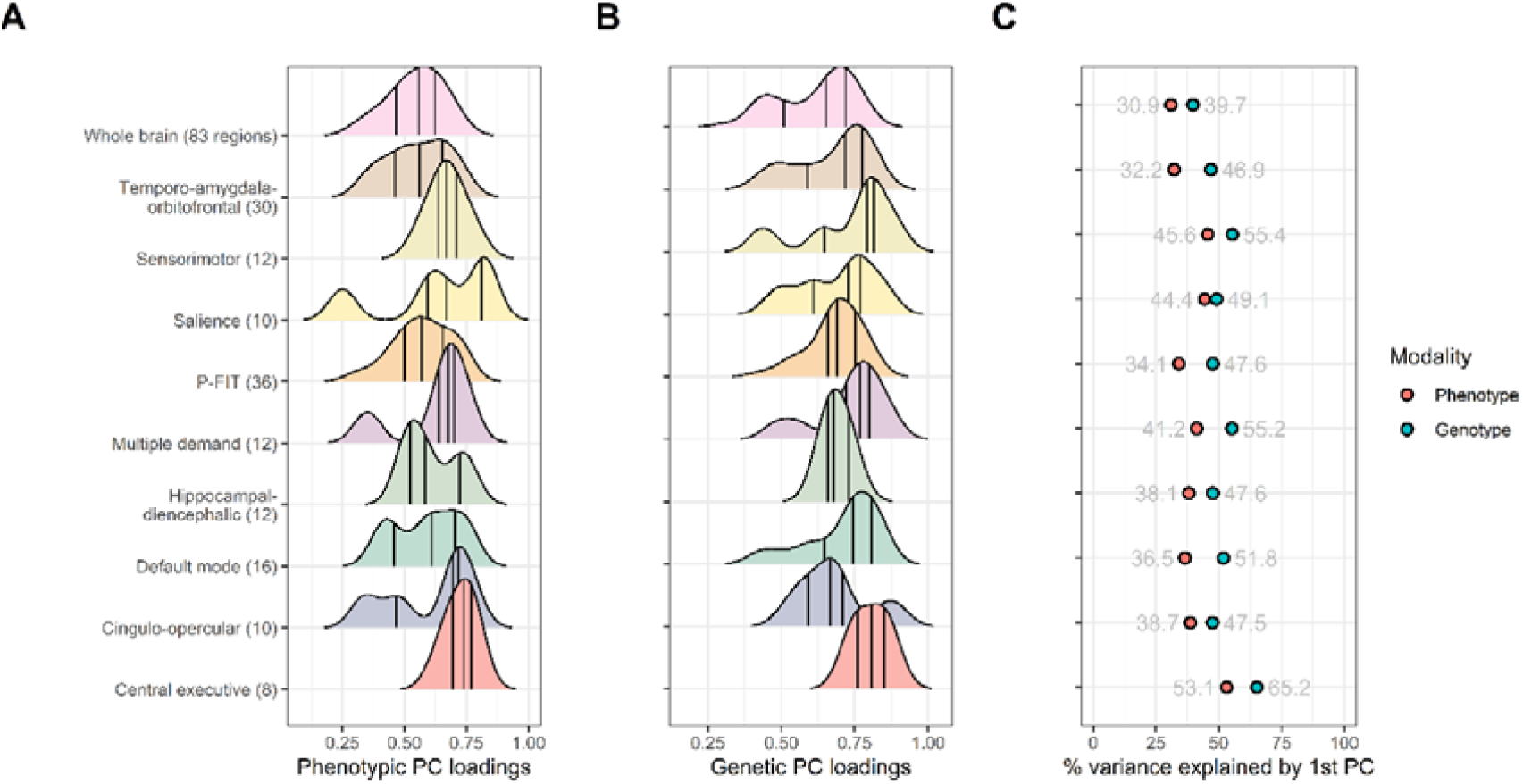
(A) Density distributions of principal component (PC) loadings on the first PC underlying phenotypic and (B) genetic networks. Vertical lines indicate quantiles. (C) Variance explained by phenotypic and genetic first PC in each network.

We used the same approach – extracting the first genetic PC and its genetic PC loadings – to examine nine predefined genetic brain-subnetworks (Fig. 1). Regions included in the networks are listed in STable 2. The percentage of genetic variance accounted for by the first network-specific PCs ranged between 65% for the central executive network and 47% for the temporo-amygdala-orbitofrontal network. While the central executive and the hippocampal-diencephalic networks had a narrow, unimodal distribution of PC loadings, the temporo-amygdala-orbitofrontal and cingulo-opercular networks had a wider, and bimodal distribution. That is, volumes included in the central executive network, for example, were more homogeneous and indexed more similar genetic variation, compared with the temporo-amygdala-orbitofrontal network (Fig. 4). Overall, percentages of explained variances were larger for networks including fewer volumes, potentially because larger networks tend to be more heterogeneous.

To test whether data-derived PCs explained more genetic variance than could be expected by chance, we present a version of Parallel Analysis which simulates PCs for uncorrelated elements with matched genetic sampling variance (see Supplementary Methods). Parallel Analysis confirmed that genetic PCs of the whole brain and the nine canonical subnetworks explained substantially more variance than expected by chance (Scree Plots SFig.s 11-20). Furthermore, we demonstrated that PCs extracted from 800 networks with randomly included brain volumes explained substantially less averaged variance than empirical canonical networks. Results from this simulation are presented in STable 5. In summary, these results illustrate that genetic dimensions of shared morphometry are well represented by the first underlying PC (i.e., accounts for the majority of genetic variance); that the dimensions differ between networks, and that they explain similar magnitudes of variance as their corresponding phenotypes.

### 3.3 General dimensions of phenotypic and genetic shared morphometry were similarly organised

To quantify how closely patterns of shared variance between phenotypic and genetic brain morphometry resemble each other, we calculated a linear regression between sets of 83 phenotypic and 83 genetic PC loadings. PC loadings indicate relative magnitudes of brain regions’ loadings on either phenotypic or genetic dimensions of shared morphometry, and serve as an index of how well a volume represents trends across the brain (or the network). The association between phenotypic PC loadings and genetic PC loadings was large and significant (*b* = 0.65, SE = 0.06, *p* = 5.07 x10^-17^, *R^2^* = 58%), indicating that an increase in one unit in the genetic PC loadings is associated with an increase of .65 units in the phenotypic PC loadings (intercept = 0.15). This approach considers ordering relative to the mean.

The Tucker congruence coefficient was used to index the degree of similarity of genetic and phenotypic PC loadings, taking into account both their relative ordering and their absolute magnitudes [34]. The Tucker coefficient revealed very high congruence in the deviation from zero between phenotypic and genetic PC loadings for the 83 volumes (Tucker coefficient = 0.99). These results illustrate a close correspondence and an equivalent organisation of phenotypic and genetic dimensions of shared morphometry; a finding that aligns with Cheverud’s Conjecture (Section 4.2).

### 3.4 Genetic dimensions of shared morphometry were associated with age sensitivity

Previous work demonstrated an association between phenotypic dimensions of shared morphometry across the whole brain, represented by phenotypic PC loadings, and indices of *age sensitivity* [3]. Age sensitivity is approximated by a correlation of a regional brain volume with age across the sample, which is typically negative in adult populations. Here, we replicated this association between phenotypic shared morphometry (i.e., phenotypic PC loadings) and age sensitivity (*r* = −0.43, *p* = 4.4 x10^-5^; Fig. 5c), and we found a significant, but smaller association for genetic PC loadings (*r* = −0.27, *p* = 0.012; Fig. 5d). This demonstrates that the more the genetic variation of a brain volume resembles general morphometric trends across the brain (larger genetic PC loading), the stronger this volume is negatively correlated with age. Note that these results emerged even though PC loadings were extracted from brain volumes *residualised* for age and were nevertheless associated with age sensitivity. In summary, these results show that phenotypic PC loadings and genetic PC loadings both display associations with age sensitivity, as indexed by cross-sectional age-volume correlations (Section 4.3).

**Fig. 5.**
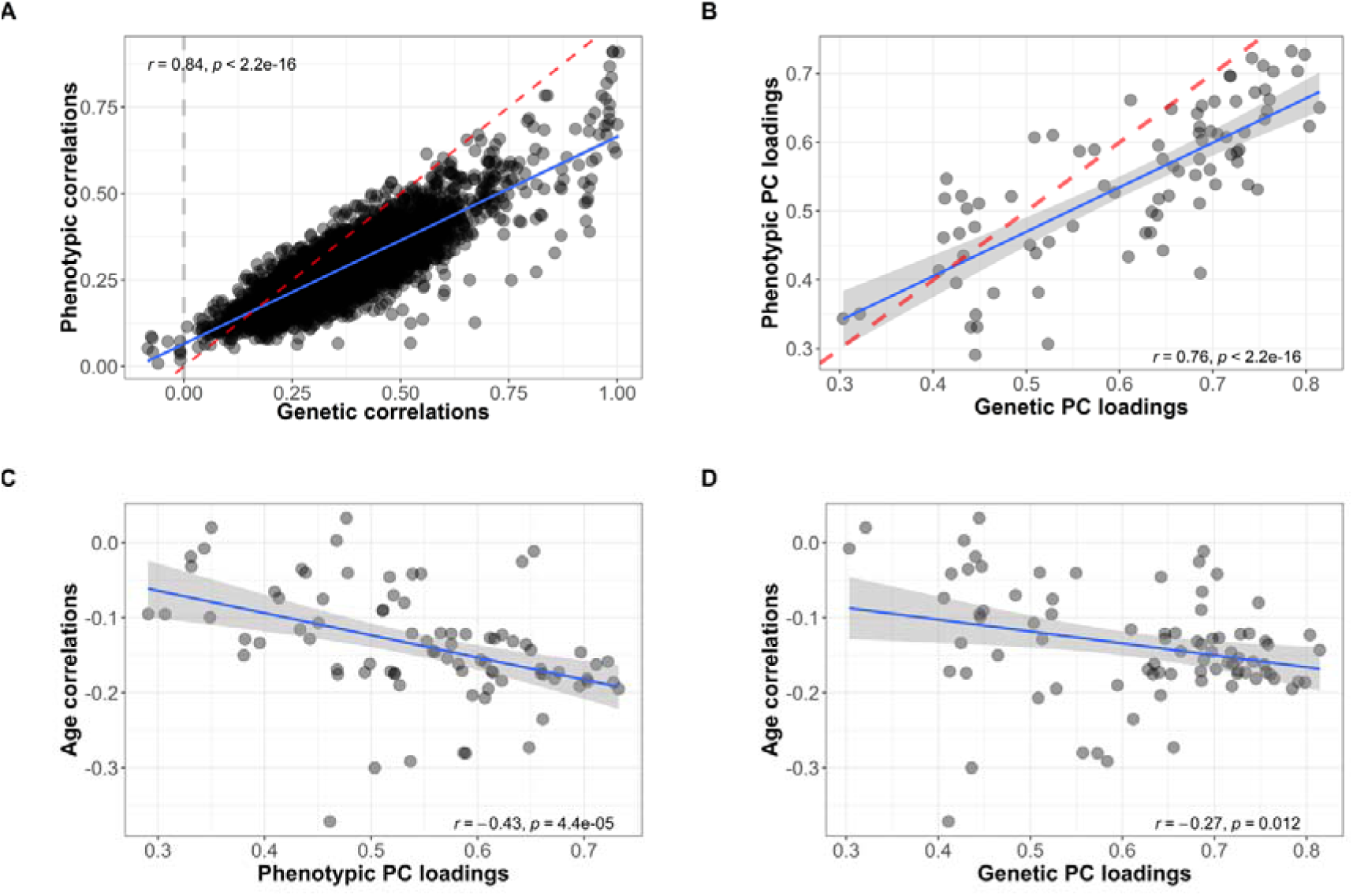
**(A)** Association between phenotypic and genetic between-region correlations of 83 regional grey-matter volumes. The dashed red line is the line of identity, with a slope of 1 and an intercept of 0. The dashed grey line indicates *r_g_* = 0. **(B)** Correlation between phenotypic and genetic PC loadings on the first PC underlying 83 regional volumes. The dashed red line is the line of identity. **(C)** Correlation between *phenotypic* PC loadings and age sensitivity as indexed by phenotypic cross-sectional age-volume correlations. **(D)** Correlation between *genetic* PC loadings and age sensitivity as indexed by phenotypic cross-sectional age-volume correlations.

### 3.5 General dimensions of shared morphometry were genetically correlated with general cognitive ability

To quantify genetic correlations between general dimensions of network morphometry and general cognitive ability, we indexed shared genetic variance across brain networks, by extracting underlying genome-wide PCs. Genome-wide PCs were calculated by summarising per-SNP effects from multiple brain volume GWAS summary statistics, weighted by volume- and network-specific PC loadings (novel method presented in Fig. 2.4). Using GenomicSEM software [32], we calculated genetic correlations between brain networks and seven cognitive traits [32] (SFig. 21). The cognitive traits mostly had high loadings on a genetic general cognitive ability factor (median = 0.81, range = 0.30-0.95); the Reaction Time task had the lowest loading on the factor (SFig. 22). Strong genetic overlap between brain networks indicated that they indexed very similar polygenic signal (*r_g_* between networks = 0.63-0.97). All networks were significantly genetically associated with the general cognitive ability factor; correlation magnitudes across all networks ranged between *r_g_* = 0.17-0.21 (Table 1). According to commonly-used rules of thumb from Hu and Bentler [35](CFI > 0.95, RMSEA < 0.08), all models showed good model fit (STable 4).

**Table 1.**
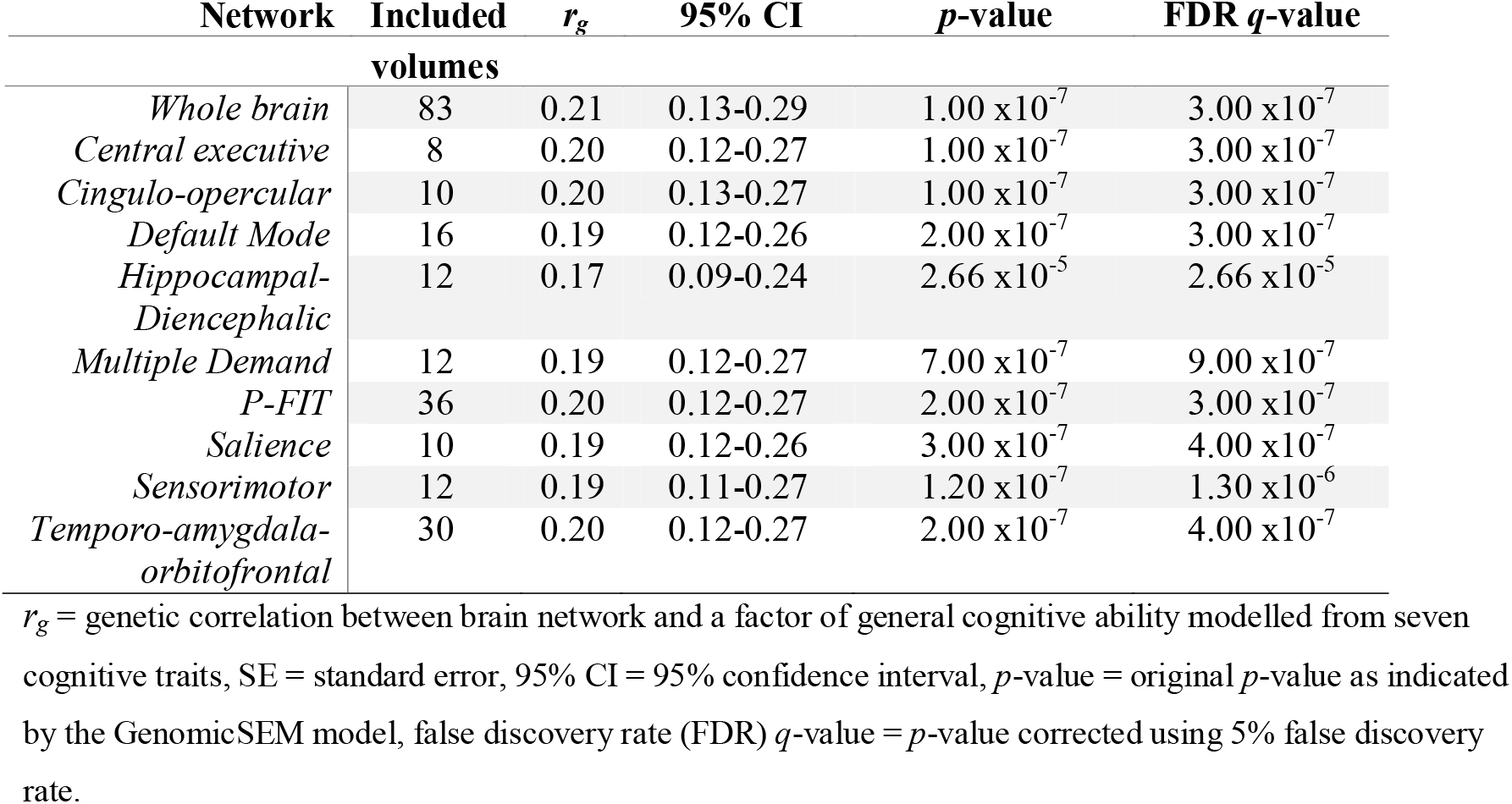
Genetic correlations between general cognitive ability and nine canonical brain networks

| Network | Included | $r_g$ | 95% CI | $p$-value | FDR $q$-value |
| --- | --- | --- | --- | --- | --- |

|  | volumes |  |  |  |  |
| --- | --- | --- | --- | --- | --- |
| *Whole brain* | 83 | 0.21 | 0.13-0.29 | 1.00 x10<sup>-7</sup> | 3.00 x10<sup>-7</sup> |
| *Central executive* | 8 | 0.20 | 0.12-0.27 | 1.00 x10<sup>-7</sup> | 3.00 x10<sup>-7</sup> |
| *Cingulo-opercular* | 10 | 0.20 | 0.13-0.27 | 1.00 x10<sup>-7</sup> | 3.00 x10<sup>-7</sup> |
| *Default Mode* | 16 | 0.19 | 0.12-0.26 | 2.00 x10<sup>-7</sup> | 3.00 x10<sup>-7</sup> |
| *Hippocampal-Diencephalic* | 12 | 0.17 | 0.09-0.24 | 2.66 x10<sup>-5</sup> | 2.66 x10<sup>-5</sup> |
| *Multiple Demand* | 12 | 0.19 | 0.12-0.27 | 7.00 x10<sup>-7</sup> | 9.00 x10<sup>-7</sup> |
| *P-FIT* | 36 | 0.20 | 0.12-0.27 | 2.00 x10<sup>-7</sup> | 3.00 x10<sup>-7</sup> |
| *Salience* | 10 | 0.19 | 0.12-0.26 | 3.00 x10<sup>-7</sup> | 4.00 x10<sup>-7</sup> |
| *Sensorimotor* | 12 | 0.19 | 0.11-0.27 | 1.20 x10<sup>-7</sup> | 1.30 x10<sup>-6</sup> |
| *Temporo-amygdala-orbitofrontal* | 30 | 0.20 | 0.12-0.27 | 2.00 x10<sup>-7</sup> | 4.00 x10<sup>-7</sup> |

$r_g$ = genetic correlation between brain network and a factor of general cognitive ability modelled from seven cognitive traits, SE = standard error, 95% CI = 95% confidence interval, $p$ -value = original $p$ -value as indicated by the GenomicSEM model, false discovery rate (FDR) $q$ -value = $p$ -value corrected using 5% false discovery rate.

Based on phenotypic findings that have highlighted the importance of the central executive network to general cognitive function [3], we hypothesised to find a stronger genetic association between general cognitive ability and the central executive network relative to other subnetworks (see pre-registered plan https://osf.io/7n4qj). There was no evidence for significant differences in total correlation magnitudes between the central executive network and general cognitive ability compared with other brain networks, even after accounting for network sizes (see Methods; Fig. 23, STable 6). We found no significant difference in model fit using GenomicSEM [32] comparing one model accounting for network size, and another model not accounting for it (Δ χ***^2^*** *p*-values ranged between .072 and 1.00; STable 5).

We also investigated whether genetic associations were driven by specific cognitive traits. We obtained non-significant *Q_trait_* heterogeneity indices [36] for all brain networks, demonstrating that the general cognitive ability factor accounted well for the patterns of association between specific cognitive abilities and the brain networks (SFig. 24). The fact that the general cognitive ability factor accounted well for specific abilities, and that the specific abilities were mostly significantly associated with the networks, confirms that the genetic associations between specific cognitive abilities and brain networks are likely general and act through a factor of general cognitive ability (Section 4.4).

### 3.6 General dimensions of shared morphometry were genetically correlated with brain age

Finally, we calculated a genetic correlation between shared morphometry across the whole brain and *brain age*. Brain age is based on individual-level predictions of how much older (or younger) an individual’s brain appears from structural MRI measures, relative to their chronological age [37] (see Supplementary Methods). We found a moderate negative genetic association (*r_g_* = −0.34; *SE* = 0.06) between general dimensions of shared morphometry across the whole-brain and brain age, suggesting that consistently larger volumes across the whole brain indicate younger brain age (Section 4.3).

## 4. Discussion

This genetically-informed study provides fundamental insights into the complex biology shared between brain organisation, ageing, and cognitive ability. Using genomic PCA, we demonstrated that general morphometric dimensions underlying brain network structures genetically overlapped with general cognitive ability, brain age, and sensitivity of a corresponding region to age, albeit being distinctly measured demographic, psychological and neuroimaging concepts. Our findings highlight measurable biological pathways giving rise to genetic variation in brain morphometry which may influence pathways underlying cognitive ability and vulnerability towards ageing. Discovery of shared genetic etiology and its associated neurodegenerative mechanisms should inform efforts of detecting and mitigating cognitive decline in ageing societies [3, 38, 39].

### 4.1 Characteristics of genetic brain network organisation

We demonstrated that genetic dimensions of shared morphometry underlying brain networks (i.e., first genetic PC) accounted for substantial systematic variance shared between brain volumes (e.g., 40% of genetic variance across the whole brain). These major genetic dimensions even explained more variance than their phenotypic analogue (31% of phenotypic variance across the whole brain). All genetic networks explained substantially more variance than was expected by chance. These findings provide a new line of evidence characterising and underpinning the existence of a genetic foundation for canonical brain networks that have featured prominently in neuroscientific studies [e.g., 7].

### 4.2 Analogous organisation of phenotypic and genetic dimensions of shared morphometry

We discovered a high degree of similarity between phenotypic and genetic features of brain network organisation (e.g., *r*_genetic vs. phenotypic correlations_ = 0.84; Tucker congruence = 0.99; *r_phenotypic vs. genetic PC loadings_* = 0.76). According to Cheverud’s Conjecture [16], this indicates that brain organisation, indexed through both phenotypic and genetic variance, seems to be underpinned by similar, overlapping developmental pathways.

### 4.3 Genetic PCs as indices of brain regions’ age sensitivity and brain age

As previously demonstrated, phenotypic PC loadings onto an underlying brain-wide dimension of shared morphometry resembled patterning of sensitivity of a corresponding region towards age (i.e., cross-sectional age-volume correlations) [3]. Here, we replicated this negative association, and showed that it also exists, albeit to a lesser degree, between age-volume correlations and genetic instead of phenotypic PC loadings. This suggests that dimensions along which brain regions share morphometric variance (i.e., generally larger volumes across an individuals’ brain) are structured similarly to patterns by which brain regions display increased vulnerability to ageing. This finding needs to be triangulated by either future longitudinal studies, or cross-sectional studies modelling within-person atrophy by incorporating information on prior brain size (e.g., intracranial volume).

One potential explanation for this association is that brain regions that are genetically predisposed to be large volumes, that share higher levels of morphometric variance with the rest of the brain, and that are more central to heavily-demanding cognitive processes, might come under more strenuous developmental and environmental pressure, perhaps through increased metabolic burden, compared with other, less central regions. Thus, the embedding of a brain volume within the whole brain’s organisation, and the genetic foundation of its positioning in the brain, could govern the functional stresses and other influences to which certain areas are exposed. This might alter disproportionately the speed at which some regions atrophy with advancing age.

That dimensions of shared morphometry resemble patterns of age sensitivity is of interest because it emerged from shared variance among brain phenotypes that had been residualised for age. Consequently, we suggest that patterns of brain structural ageing, a construct labelled *brain age* [29], might not capture how quickly an individual’s regional volumes decline compared to their peers, but rather, general healthy morphometry across the brain. Previous research showed that a younger-appearing brain, relative to the individual’s chronological age, predicted better physical fitness, better fluid intelligence, and longevity [29]. Healthy brain morphometry could vary between people for many non-age-related reasons, including genetic predisposition. Individuals that are genetically predisposed towards consistently larger brain volumes might have generally healthier, better-integrated brains, which could be more resilient towards harmful environmental factors.

In line with this theory, we found that younger brain age was genetically associated with a major dimension of brain-wide shared morphometry as indexed by a genetic PC (*r_g_* = - 0.34; *SE* = 0.06). Thus, consistently larger volumes across the brain indicate a younger structural brain organisation, and this is the first study to quantify the degree to which these two concepts overlap. It motivates further investigation into the possibility that they are underpinned by the same general shared biological pathways.

### 4.4 Genetic PCs as indices of cognitive performance

This study demonstrated that cognitive ability is positively associated with genetic morphometric variance shared across the whole brain, and across smaller canonical networks. This was investigated by modelling a genetic factor of general cognitive ability using GenomicSEM [32]. We calculated the genetic correlation between general cognitive ability and genetic PCs across the whole brain, and nine canonical subnetworks. The whole brain and all nine networks were significantly genetically correlated with general cognitive ability at magnitudes between 0.17 and 0.21. This was the same level of genetic association with general cognitive ability that was previously found for broad measures of total brain volume [40]. There was no evidence to suggest that those magnitudes statistically differed between the networks; probably because the polygenic signal indexed by the genetic PCs were highly similar between brain networks (mean *r_g_* between networks 0.83, SD = 0.09).

This indicates that the genetic association between brain morphometry and cognitive ability was not driven by specific network configurations. Instead, genetic PCs indexed genetic variance relevant to larger brain volumes and a brain organisation that is advantageous for better cognitive performance. This was regardless of how many brain regions and from which regions the measure of shared genetic morphometry was extracted. This lack of differentiation between networks, in how strongly they correlate with cognitive ability, is in line with the suggestion that the total number of neurons in the mammalian cortex, which should at least partly correspond to its volume, is a major predictor of higher cognitive ability [41]. These findings suggest that highly shared brain morphometry between regions, and its genetic analogue, predict a generally bigger, and cognitively better-functioning brain.

Unexpectedly, genetic correlations between networks and cognitive ability did not suggest any prominent role of the central executive network (a previous *phenotypic* study [3] demonstrated that the central executive network was disproportionately predictive of cognitive abilities relative to its few included volumes). On a genetic level of analysis, we also expected a stronger correlation with cognitive ability for the central executive network compared with the other networks. The lack of differentiation between networks, taken together with previous phenotypic evidence for a disproportionately large association between the cognitive ability and the central executive, suggests nongenetic mechanisms to play important roles, perhaps developmental and environmental influences, through which the central executive network matures, and specialises for cognitive performance.

### 4.5 Limitations

Analyses in this study come with limitations. Genetic correlations are representative for genetic associations across the entire genome, but do not give direct insight into specific genomic regions of sharing. As genetic correlations were calculated using LDSC, the limitations that apply to LDSC methodology are relevant to our study (discussion in Supplementary Note). We conclude based on heritability estimates, indexing signal-to-noise ratios in GWAS, that there was sufficient polygenic signal to warrant LDSC analysis (heritability ranged 7-42%). LDSC intercepts were perfectly associated with phenotypic correlations (*R^2^* = 0.99), indicating that the analyses successfully separated confounding signal (including environmental factors) from the estimates of genetic correlations.

This study was conducted in the UK Biobank sample, which is not fully representative of the general population: its participants are more wealthy, healthy and educated than average [42]. Cohort effects may affect the degree to which differential cortical regional susceptibility to ageing can be inferred from cross-sectional data. It remains to be tested whether our results can be extrapolated to socio-economically poorer subpopulations, or outside European ancestry. Results were also dependent on the choice of brain parcellation to divide the cortex into separate regions.

### 4.6 Conclusion

This genetically-informed study delivered evidence for shared etiology between factors that may contribute to neurodegenerative mechanisms underlying ageing-related cognitive decline. Using genome-wide data, we quantified a substantial overlap of genetic variation between distinct measures of ageing, cognitive ability, and brain morphometry, all of which are variables of interest due to their potential social and economic consequences for ageing societies. These fundamental insights will help guide investigations into risk factors, biological mechanisms, and potential interventions of ageing-related cognitive decline.

More specifically, we demonstrated that younger brain age genetically captured interindividual variation substantially related to brain network structures (i.e., consistently enlarged volumes). Because the network structures were modelled based on variance independent of age, this suggests that younger brain age could primarily be an index of brain health. Contrary to previous phenotypic findings, our genetic analyses did not provide evidence for a disproportionate role of the central executive network in cognitive performance. This motivates future investigations into environmental influences on the specialisation of brain networks. Altogether, our new genomic PCA methodology and the resulting insights of this study provide a basis for future investigations that aim to interrogate the genetic and environmental bases of ageing and cognitive decline.

## Supplementary Methods

### Study Design

#### UK Biobank data

Magnetic resonance imaging (MRI) data was collected by the UK Biobank study with identical hardware and software in Manchester, Newcastle, and Reading. Brain volumetric phenotypes were pre-processed by an imaging-pipeline developed and executed on behalf of UK Biobank [43]. More information on T1 processing can be found in the UK Biobank online documentation [44]. Briefly, cortical surfaces were modelled using FreeSurfer, and volumes were extracted based on Desikan-Killiany surface templates [45]; subcortical areas were derived using FreeSurfer aeseg tools [46]. Volumetric measures (mm^3^) have been generated in each participant’s native space. We used 83 available imaging-derived phenotypes (IDPs) of cortical and subcortical grey-matter volumes in regions of interest spanning the whole brain (UK Biobank category 192 & 190; STable 1). We assume the IDPs to be normally-distributed.

#### Phenotypic quality control

Excluding participants who withdrew consent, we considered 41,776 participants with non-missing T1-weighted IDPs that had been processed in conjunction with T2-weighted FLAIR (UK Biobank field ID 26500) where available. Using both T1 and T2 measures ensures more precise cortical segmentation [47]. Extreme outliers outside of 4 standard deviations from the mean were excluded, which resulted in between 41,686 to 41,769 available participants depending on the IDP. 381 participants were excluded as they self-reported non-European ethnicity. Across the 83 brain volumes variables and the covariates, this phenotypic quality control resulted in 39,947 complete cases, for whom the following genetic quality control steps were performed.

#### Genetic quality control

Out of the 39,947 UK Biobank participants, genetic data were available for 38,957 participants. Genetic data was quality controlled on by UK Biobank and were downloaded from the full release [48]. We applied additional quality control as previously described in Coleman et al. [49] using PLINK2 [50]. 38,038 participants were of European ancestry according to 4-means clustering on the first two genetic principal components available through UK Biobank [51]. Of those participants, we removed 72 due to quality assurance provided by UK Biobank and 204 participants due to high rates of missingness (2% missingness). To obtain a sample of unrelated individuals, 956 participants were removed using the greedyRelated algorithm (KING *r* < 0.044 [52]). The algorithm is “greedy” because it maximises sample size; for example, it removes the child in a parent-child-trio. Finally, 28 participants were removed because genetic sex did not align with self-reported sex, resulting in a total of 36,778 participants (STable 10). Genetic sex was identified based on measures of X-chromosome homozygosity (*F*_X_; removal of participants with *F*_X_⎕<⎕0.9 for phenotypic males, *F*_X_⎕>⎕0.5 for phenotypic females). The final sample (*N* = 36,778) included 19,888 females (54 %) and had an average age of 63.3 years at the neuroimaging visit (range from 40.0 to 81.8 years).

Out of 805,426 available directly genotyped variants, 104,771 were removed for high rates of missing genotype data (> 98%). 103,137 variants were removed due to a minimum allele frequency of 0.01, and 9,935 variants were removed as they failed the Hardy-Weinberg exact test (*p*-value = 10^-8^). After excluding 16,326 variants on the sex chromosomes and those with chromosome labels larger than 22, we obtained a final sample of 571,257 directly genotyped SNPs. Imputed genotype data was obtained by UK Biobank with reference to the Haplotype Reference Consortium [53], and we filtered them for a minor allele frequency of above 0.01 and an IMPUTE INFO metric of above 0.4.

#### Measures of cognitive performance

UK Biobank collected cognitive performance data using assessment on a touchscreen computer. The following seven tests were implemented: *Matrix Pattern Completion task* for nonverbal reasoning, *Memory – Pairs Matching Test* for memory, *Reaction Time* for perceptual motor speed, *Symbol Digit Substitution Task* for information processing speed, *Trail Making Test – B* and *Tower Rearranging Task* for executive functioning, and *Verbal Numerical Reasoning Test* for verbal and numeric problem solving, or fluid intelligence. Despite the non-standard and unsupervised delivery of assessment, these cognitive tests demonstrate strong concurrent validity compared with standard reference tests (*r* = .83) and good test-retest reliability (Pearson *r* range for different cognitive tests = 0.4–0.78) [54].

In this study, we considered GWAS summary statistics of performance in these seven cognitive tests by de la Fuente, Davies [12] that were calculated with between 11,263 and 331,679 participants for each test. We consider the HapMap 3 reference SNPs with the MHC regions removed.

### Statistical analysis

#### GWAS summary statistics calculation

GWAS summary statistics for the 83 regional brain volumes (continuous variables) were calculated using REGENIE [31], which fits polygenic effects in a linear mixed model using Ridge regression. The REGENIE pipeline is split into two steps: First, blocks of directly genotyped SNPs are used to fit a cross-validated whole-genome regression model using Ridge regression, to determine the amount of phenotypic variance explained by genetic effects. Second, the association between the phenotype and imputed genetic variants is calculated conditional upon Ridge regression predictions from the first step. Proximal contamination is circumvented by using a leave-one-chromosome-out scheme.

Covariates included in the GWAS analyses were *age at neuroimaging visit*, *sex*, *genotyping batch*, and *40 genetic principal components* as provided by UK Biobank. We also derived the variables *time of year*, *head position*, and *acquisition site,* but excluded them from our set of GWAS covariates because they were not associated with the brain volumes at the pre-registered arbitrary cut-off of *r* ≤ .10 (STable 9), and therefore explained less than 1% of the phenotype variance. Note that, in contrast to other existing brain-volume GWAS in UK Biobank [e.g., 55], our analyses were conducted *without* controlling for brain size (or any other global brain measure such as total grey-matter volume or intracranial volume). Genetic correlations calculated relative to such global measures are known to attenuate genetic correlations among volumes, as well as with other traits such as cognitive abilities [15]. In the context of this study, we aim to model general dimensions of variance shared between brain volumes which will closely covary with brain size. Attenuated genetic correlations would hide major dimensions of variance across genetic brain networks, because much of the variance shared between volumes overlaps with variance indexed by brain size and would therefore not tag general dimensions of shared genetic variance between brain volumes. This variance is of interest because general intelligence yields global rather than a region-specific associations with grey matter volume [28]. Equally, aging affects the whole brain rather than individual regions [56].

#### Genetic and phenotypic correlation matrices between brain volumes

To derive dimensions of shared morphometry across brain volumes, we calculated both a phenotypic and a genetic correlation matrix from 83 grey-matter volume variables. Phenotypic regional brain volumes were residualised for age at neuroimaging visit and sex, and then used to estimate a phenotypic correlation matrix through Pearson’s correlations with complete pairwise observations. The genetic correlation matrix was inferred through LDSC, a technique quantifying shared polygenic effects between traits using GWAS summary statistics. Cross-trait LDSC regresses the product of effect sizes in two GWAS onto linkage disequilibrium scores, indicating how correlated a genetic variant is with its neighbouring variants [33]. The slope indexes the genetic correlation, while the intercept captures signal uncorrelated with LD, such as population stratification, environmental confounding, and sample overlap.

To quantify the relationship between phenotypic and genetic correlations, we estimated the correlation between 3403 phenotypic and genetic between-region correlations (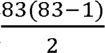 = 3403 correlations between 83 volumes). Additionally, we calculated genetic correlation matrices for smaller canonical networks including fewer brain volumes than the whole brain. For example, the central executive network included eight regional volumes (STable 2 lists volumes included in the nine canonical networks). We reported SNP-heritability estimates for each brain volume inferred through LDSC.

#### Principal component analysis (PCA) of genetic and phenotypic correlation matrices

PCA was applied to the phenotypic and genetic correlation matrices indicating genetic overlap between brain volumes described above to obtain their respective first principal component (PC). The first PC represents an underlying dimension of common structural sharing across regional volumes, which we refer to as general dimensions of shared morphometry throughout this manuscript. PC loadings were calculated for all volumes in the whole brain, as well as volumes in smaller canonical networks to quantify contributions of regional volumes to this either brain-wide, or network-specific dimension of shared morphometry.

#### Parallel analysis

We tested whether genetic PCs explained more variance than expected by chance, that is, whether they explained more than 95% of their corresponding PCs generated under a simulated null correlation matrix. We developed a version of parallel analysis to generate null distributions of eigenvalues by simulating null correlation matrices sampled from a diagonal population correlation matrix, where the multivariate sampling distribution is specified to take the form of the sampling distribution of the standardised empirical genetic correlation matrix (the V_STD_ matrix, as estimated using GenomicSEM [32]). This sampling correlation matrix serves as an index of the precision of the elements in the empirical genetic covariance matrix (i.e., heritabilities and co-heritabilities across traits) and the sampling dependencies among these when generating the random null models. We specified 1,000 replications to simulate the null correlation matrices and use a 95% threshold for distinguishing true eigenvalues from noise.

#### Simulation of networks with randomly included brain volumes

We performed an additional sensitivity analysis simulating networks with randomly included brain volumes, to determine whether shared structural variance relied on network membership, or arose through phenotypic properties common to all regional brain volumes. To compare explained variances between canonical networks and random networks, we quantified the expected explained variance in random networks by randomly sampling regions 800 times each, for different numbers of included volumes (because networks including fewer volumes generally tend to explain a larger percentage of variance, as larger networks are more heterogeneous). That is, simulations were run for 8, 10, 12, 16, 30, and 36 included regions, to obtain a distribution for each networks size to compare the corresponding network’s explained variance to. We reported the mean explained variance by PCs for networks with randomly included volumes and a 95% confidence interval. Comparisons between explained variances for random and empirical networks were done for the same number of included volumes.

#### Correlation between phenotypic and genetic PC loadings

To compare whether genetic correlations structures of regional brain morphometry resembled the phenotypic correlation structure of the same regions, we calculated an un-standardised linear regression with a vector of 83 phenotypic whole-brain PC loadings as the dependent variable, and a vector containing 83 genetic whole-brain PC loadings as the independent variable. We calculated the Tucker congruence coefficient to quantify the relative similarity between the two sets of PC loadings independent of their absolute magnitude. The coefficient is insensitive to scalar multiplication [57].

#### Correlation between genetic PC loadings with age sensitivity

Pearson’s correlations between 83 phenotypic grey-matter volumes and age at neuroimaging visit were calculated to quantify cross-sectional age-volume-correlations for each of the 83 brain volumes. These age-volume correlations are referred to as *age sensitivity* throughout the rest of the manuscript. We estimated the correlation between a vector containing indices of age sensitivity and (1) a vector of *genetic* whole-brain PC loadings, and for comparison (2) a vector of *phenotypic* whole-brain PC loadings.

#### Genome-wide shared genetic variance of morphometry across the whole brain and canonical networks

To statistically represent genome-wide shared morphometric variance across brain volumes (i.e., genetic PCs), we developed a novel method summarising genome-wide by-variant effects contained in the grey-matter volume GWAS summary statistics, which were weighted by their respective (region-specific) PC loadings obtained through PCA. We derived GWAS summary statistics for a genetic principal component of multiple GWAS phenotypes derived from samples of unknown degrees of overlap by adapting existing software for genome-wide multivariate meta-analysis by Baselmans et al. [58] and using GenomicSEM [32]. Fig. 2 illustrates this approach in a four-step procedure. The input data for our approach are GWAS summary statistics for 83 cortical and subcortical brain volumes (step 1). We have made them publicly available online. Using the GenomicSEM software [32], we obtained a genetic correlation matrix indicating genetic overlap between these 83 brain volumes (step 2). We extracted PC loadings on the underlying general dimension of shared genetic variance for each of the 83 regions (step 3). Finally, we modified the existing genome-wide multivariate meta-analysis software package [58], in order to create summary statistics for an underlying genetic PC. Genome-wide SNP effects were calculated as an average of all SNP effects contributed by the 83 GWAS phenotypes, weighted by their respective PC loading, with standard errors computed using a method that corrects for sample overlap, as estimated by LDSC (step 4). We used this approach to calculate univariate summary statistics to represent general dimensions of shared morphometry between regional volumes across the whole brain (83 GWAS phenotypes), as well as nine smaller canonical networks.

We had tested and validated this novel approach in an independent set of GWAS summary statistics of four risky behaviours [59]. In addition to the risky behaviour GWAS, another set of summary statistics is available for a phenotypic PC underlying these risky behaviour phenotypes that the authors had calculated phenotypically before running GWAS analyses. We compared these phenotypic PC GWAS summary statistics by Linnér, Biroli [58] with summary statistics for a *genetic* PC underlying the four risky behaviours GWAS that we calculated using our novel method outlined above (Fig. 2). We found that they correlated at a magnitude of *r_g_* = 0.99 (*SE* = 0.037) confirming that our method captures the same signal as can be obtained from phenotypic PCs, by simply relying on publicly available GWAS data. For details of the analysis and code refer to: https://annafurtjes.github.io/genomicPCA/.

#### Genetic correlation between general dimensions of shared morphometry across the whole-brain and brain age

Using LDSC [33], we calculated a genetic correlation between genetic morphometric sharing across the whole brain and *brain age*. The summary statistics indexing dimensions of shared morphometry across brain volumes were created using the novel method presented above (Fig. 2). We downloaded the brain age GWAS summary statistics online [37]. Brain age is a phenotype based on individual-level predictions of how much older (or younger) an individual’s brain appears, relative to their chronological age. It is estimated using parameters characterising the relationship between age and structural neuroimaging measures (volume, thickness, and surface area) that were tuned using machine learning in an independent sample. The final brain age phenotype indexed in the GWAS was calculated as the difference between participants chronological age and their age as predicted based on structural brain characteristics.

#### Genetic correlations between brain networks and a factor of general cognitive ability

We assessed genetic correlations between brain networks and general cognitive ability using GenomicSEM [32]. Using univariate network-specific summary statistics (as describe above; Fig. 2) and a genetic general cognitive ability factor modelled from seven cognitive ability GWAS summary statistics, the GenomicSEM software [32] was used to model general cognitive ability and perform multivariate LDSC using diagonally weighted least squares. To quantify model fit, we reported default fit indices calculated by the GenomicSEM package: χ*^2^* values, the Akaike Information Criterion (AIC), the Comparative Fit Index (CFI) and the Standardised Root Mean Square Residuals (SRMR). The multiple testing burden was addressed by correcting *p*-values from the genetic correlations for multiple testing with a false-positive discovery rate of 5% [60].

We preregistered that we would test for significant differences in correlation magnitudes between the networks that yielded a significant association with general cognitive abilities. Because we hypothesised a particularly strong association for the central executive network, we planned to perform this comparison between the central executive and all other networks, to reduce the multiple testing burden. We fitted two GenomicSEM models in which correlation magnitudes between general cognitive ability and both the central executive and another network were either freely estimated, or they were forced to be the same. A significant decrease in model fit between the freely estimated model and the constrained model (*df* = 1) would indicate that there likely are differences in correlation magnitudes between the networks in how strongly they correlate with general cognitive ability (SFig. 23).

Additionally, we assessed whether the central executive network was disproportionately genetically correlated with general cognitive ability considering its small size (i.e., few included volumes). Similar to the approach described above, we fitted two models: One, in which we freely estimate the correlation between the central executive and general cognitive ability, and the correlation between another network and general cognitive ability. We then divided the correlation magnitude by the number of regions included in the network (i.e., magnitude was divided by 8 for the central executive network, it was divided by 16 for the default mode, by 36 for the P-FIT etc.). The second model had the same set up, but we forced the adjusted correlations for the two networks to be equal (e.g., r_central executive_ / 8 == r_default_ / 16). We assessed whether there was a significant difference in χ***^2^*** model fit between these two models. As above, a significant decrease in model fit between the freely estimated model and the constrained model (*df* = 1) would indicate that there likely are differences in relative correlation magnitudes (i.e., magnitudes adjusted for network sizes). Based on previous findings, we expected the relative magnitude for the central executive network to be significantly larger than the relative magnitude for any other network.

To probe whether any specific cognitive ability might have driven the genetic associations between brain networks and general cognitive ability, we reported genetic correlations between the significant networks and three specific cognitive abilities: (1) *Matrix Pattern Completion task* to represent nonverbal reasoning, (2) *Memory – Pairs Matching Test* to represent memory, and (3) *Symbol Digit Substitution Task* to represent information processing speed. Reducing the analyses to only three consistent and representative cognitive measures reduced the burden of multiple testing.

We calculated *Q_trait_* heterogeneity indices [36] to evaluate whether the general cognitive ability factor that we fit in the models above accounts well for the specific cognitive abilities. To this end, we compared the fit of two models for each network as displayed in SFig. 24. One model allows for independent associations between the seven cognitive traits, and both general cognitive ability and the brain network. The second model forces the association between the seven cognitive traits and the brain network to go through the general cognitive ability factor. We obtained χ***^2^*** fit statistics for both models and tested their difference for statistical significance (Δ χ***^2^*** ≠ 0; *df* = 6). Non-significant results (*p* > 0.05/10) would suggest that genetic associations between cognitive abilities and brain networks are likely general and act through a factor of general cognitive ability.

#### Data and code availability

Access to phenotypic and genetic UK Biobank data was granted through the approved application 18177. We have made the 83 GWAS summary statistics of regional volumes available at the GWAS catalogue (https://www.ebi.ac.uk/gwas/). GWAS summary statistics for the seven cognitive traits by de la Fuente, Davies [12] were downloaded at https://datashare.ed.ac.uk/handle/10283/3756. The pre-registration for this analysis can be found online (https://osf.io/7n4qj). Full analysis code including results for this study are available at https://annafurtjes.github.io/Genetic_networks_project/index.html.

## Acknowledgements

AEF is funded by the Social, Genetic and Developmental Psychiatry Centre, King’s College London and the National Institute of Health (NIH) grant R01AG054628. SJR is funded by the Jacobs Foundation. JHC is funded by a UK Research & Innovation(UKRI) Innovation Fellowship (MR/R024790/1; MR/R024790/2). JF is funded by the National Institutes of Health (NIH) grant R01AG054628. JF and EMTD are members of the Population Research Center (PRC) and Center on Aging and Population Sciences (CAPS) at The University of Texas at Austin, which are supported by NIH grants P2CHD042849 and P30AG066614. JD, JWM, and EMTD were supported by NIH R01AG054628. IJD is with the Lothian Birth Cohorts group, which is funded by Age UK (Disconnected Mind grant), the Medical Research Council (grant no. MR/R024065/1) and the University of Edinburgh’s School of Philosophy, Psychology and Language Sciences. The contribution by RA represents independent research part-funded by the National Institute for Health Research (NIHR) Maudsley Biomedical Research Centre at South London and Maudsley NHS Foundation Trust and King’s College London. The views expressed are those of the author(s) and not necessarily those of the NHS, the NIHR or the Department of Health and Social Care. The contribution by JRIC represents independent research part-funded by the National Institute for Health Research (NIHR) Maudsley Biomedical Research Centre at South London and Maudsley NHS Foundation Trust and King’s College London. The views expressed are those of the authors and not necessarily those of the NHS, the NIHR or the Department of Health and Social Care. SRC is supported by a Sir Henry Dale Fellowship jointly funded by the Wellcome Trust and the Royal Society (Grant Number 221890/Z/20/Z). This research was funded in part by the Wellcome Trust [221890/Z/20/Z]. For the purpose of open access, the author has applied a CC BY public copyright licence to any Author Accepted Manuscript version arising from this submission.

The authors gratefully acknowledge the UK Biobank resource (https://www.ukbiobank.ac.uk/) and its research team, who have made this work possible (project number 18177). The authors acknowledge use of the research computing facility at King’s College London, Rosalind (https://rosalind.kcl.ac.uk), which is delivered in partnership with the National Institute for Health Research (NIHR) Biomedical Research Centres at South London & Maudsley and Guy’s & St. Thomas’ NHS Foundation Trusts, and part-funded by capital equipment grants from the Maudsley Charity (award 980) and Guy’s & St. Thomas’ Charity (TR130505).

## Author contributions

Conceptualisation and methodology: SJR, EMTD, JHC, AEF, SRC

Supervision: SJR, EMTD, JHC

Network characterisation: SRC

Idea to investigate genetic brain age – shared morphometry correlation: JWM

Script used to perform genetic parallel analysis: JF

Data access: CML

Genetic quality control: AEF, JRIC

GWAS calculation: AEF, RA

Data analysis: AEF

Writing: AEF

Visualisations: AEF

Reviewed draft: all authors

## Supplemental Information titles and legends

*Supplementary Table 1*. 83 cortical and subcortical grey-matter regions of interest

*Supplementary Table 2*. Network characterisation

*Supplementary Table 3*. Explained variance and descriptive statistics of PC loadings within phenotypic canonical networks

*Supplementary Table 4*. Model fit for genetic correlations between genetic general cognitive ability and each canonical network

*Supplementary Table 5*. Fit indices for the comparison between freely-varying or constrained correlations with general cognitive ability between central executive and other networks

*Supplementary Table 6*. Fit indices for the *adjusted* comparison between freely-varying or constrained correlations with general cognitive ability between central executive and other networks

*Supplementary Table 7*. Genetic correlations between three cognitive abilities and brain networks

*Supplementary Table 8*. Canonical networks explain more variance than networks with randomly included volumes

*Supplementary Table 9*. Associations between brain volumes and potential covariates

*Supplementary Table 10*. Genetic quality control exclusion criteria resulting in a total GWAS sample of 36,778 out of 39,947 participants

*Supplementary Fig. 1*. Genetic correlation matrix inferred through LDSC across the whole brain (83 volumes).

*Supplementary Fig. 2*. Genetic correlations inferred through LDSC among the central executive network (8 volumes).

*Supplementary Fig. 3*. Genetic correlations inferred through LDSC among the cingulo-opercular network (10 volumes).

*Supplementary Fig. 4*. Genetic correlations inferred through LDSC among the default mode network (16 volumes).

*Supplementary Fig. 5*. Genetic correlations inferred through LDSC among the hippocampal-diencephalic network (12 volumes).

*Supplementary Fig. 6*. Genetic correlations inferred through LDSC among the multiple demand network (12 volumes).

*Supplementary Fig. 7*. Genetic correlations inferred through LDSC among the P-FIT network (36 volumes).

*Supplementary Fig. 8*. Genetic correlations inferred through LDSC among the salience network (10 volumes).

*Supplementary Fig. 9*. Genetic correlations inferred through LDSC among the sensorimotor network (12 volumes).

*Supplementary Fig. 10.* Genetic correlations inferred through LDSC among the temporo-amygdala-orbitofrontal network (30 volumes).

*Supplementary Fig. 11.* Parallel analysis in the central executive network

*Supplementary Fig. 12.* Parallel analysis in the cingulo-operular network

*Supplementary Fig. 13.* Parallel analysis in the default mode network

*Supplementary Fig. 14.* Parallel analysis in the hippocampal-diencephalic network

*Supplementary Fig. 15.* Parallel analysis in the multiple demand network

*Supplementary Fig. 16.* Parallel analysis in the P-FIT network

*Supplementary Fig. 17.* Parallel analysis in the salience network

*Supplementary Fig. 18.* Parallel analysis in the sensorimotor network

*Supplementary Fig. 19.* Parallel analysis in the temporo-amygdala-orbitofrontal network

*Supplementary Fig. 20.* Parallel analysis in the whole brain with 83 nodes

*Supplementary Fig. 21.* Genetic correlations between seven cognitive traits and brain networks. Descriptively, performance in the Tower Rearranging Task has the largest association with brain networks in comparison with other cognitive tasks. Abbreviations: Matrix = Matrix Pattern Completion task; Memory = Memory – Pairs Matching Test; RT = Reaction Time; Symbol Digit = Symbol Digit Substitution Task; Trails-B = Trail Making Test – B; Tower = Tower Rearranging Task; VNR = Verbal Numerical Reasoning Test; central exec = central executive; cingulo = cingulo-opercular; hippocampal = hippocampal-diencephalic; multiple = multiple demand; p fit = parieto-frontal integration theory; sensori = sensorimotor; temporo = temporo-amygdala-orbitofrontal

*Supplementary Fig. 22.* Genetic correlation between the central executive network and factor *g* modelled for correlation structure of seven cognitive traits. The seven cognitive traits and the network are inferred through LDSC, and the factor through factor analysis. Matrix = Matrix Pattern Completion task; Memory = Memory – Pairs Matching Test; RT = Reaction Time; Symbol Digit = Symbol Digit Substitution Task; Trails-B = Trail Making Test – B; Tower = Tower Rearranging Task; VNR = Verbal Numerical Reasoning Test. Model fit: χ*^2^* = 124.04, df = 20, *p*-value = 2.1 x10^-20^, AIC = 174.04, CFI = 0.97, SRMR = 0.079

*Supplementary Fig. 23.* Illustration of the genomic structural equation models used to test whether correlation magnitudes with genetic general cognitive ability differ between the central executive network and other significantly associated networks. The model on the right freely estimates correlation parameters between two networks and genetic g while allowing for correlations between the networks. In the left model, we force the correlation magnitudes to be the same, and assess whether model fit deteriorates significantly, to conclude whether correlation magnitudes between networks are likely different from each other.

*Supplementary Fig. 24.* Structural equation models to calculate *Q_trait_* heterogeneity indices

