## Supplementary_Figures for "General dimensions of human brain morphometry inferred from genome-wide association data"

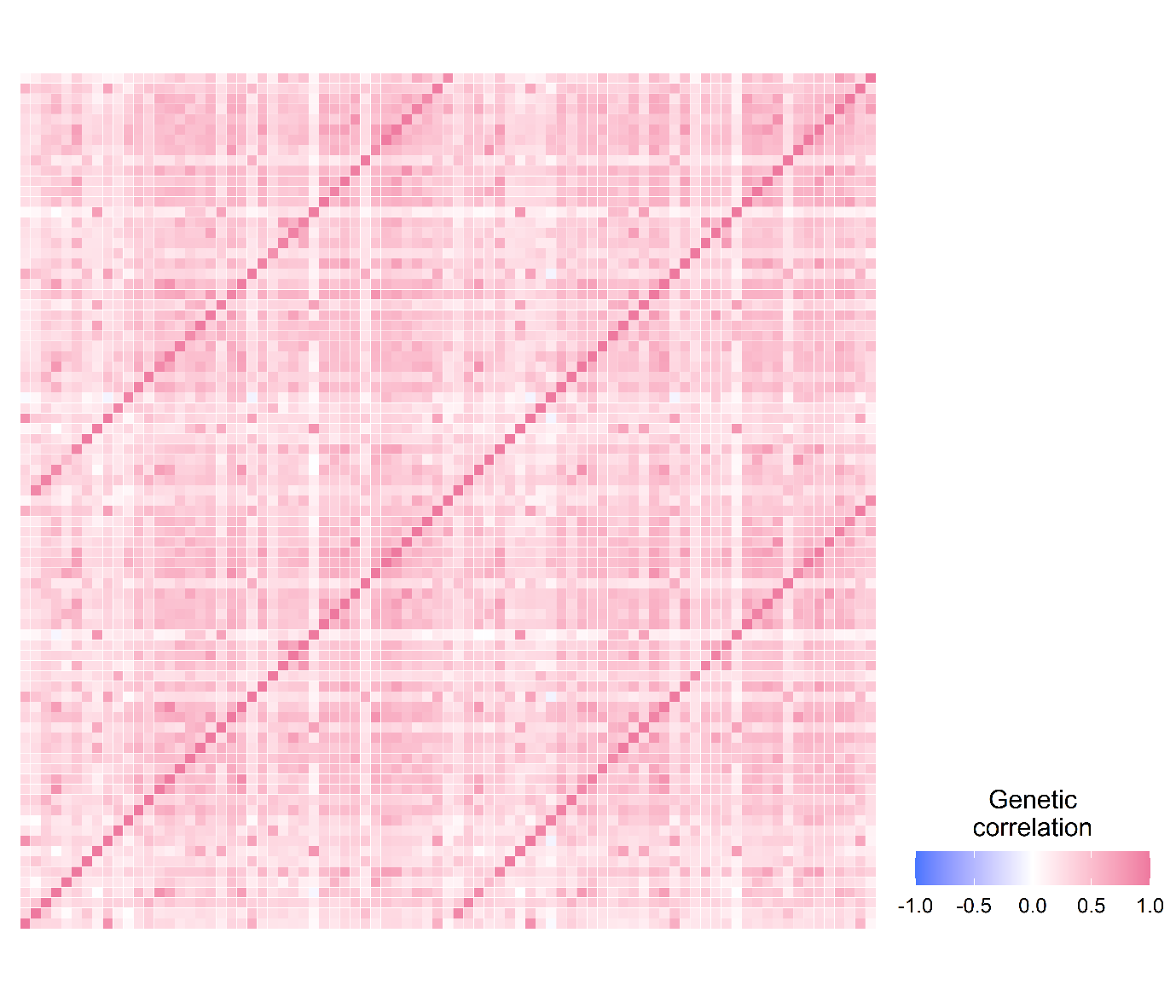

*Supplementary Figure 1.* Genetic correlation matrix inferred through bivariate LDSC across the whole brain (83 volumes). High correlations along the off-diagonal lines, parallel to diagonal, indicate nearly perfect correlations between regions and their homologous counterparts in the opposite hemisphere (brain stem is an exception as it has no counterpart in the opposite hemisphere).

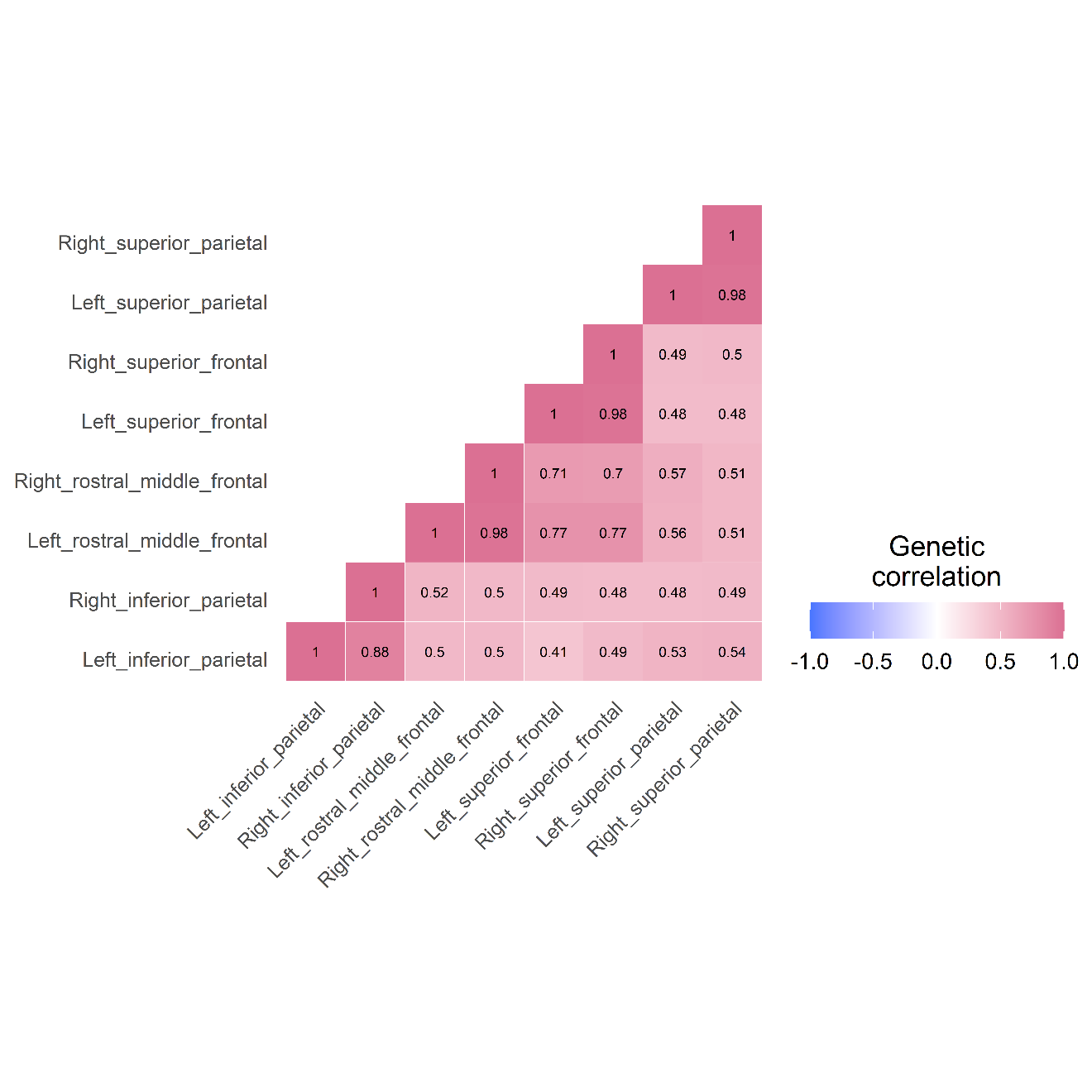

*Supplementary Figure 2.* Genetic correlations inferred through LDSC among the central executive network (8 volumes).

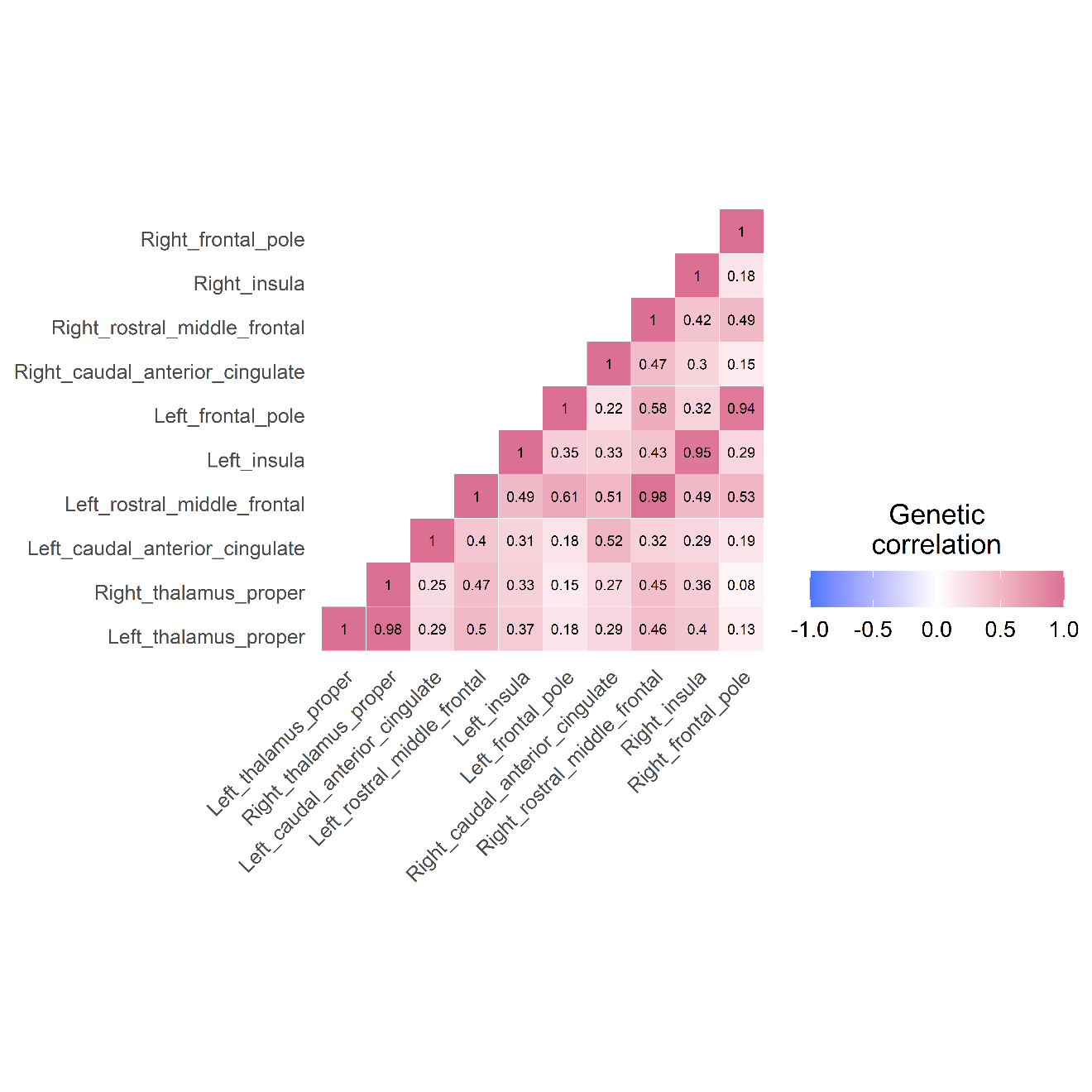

*Supplementary Figure 3.* Genetic correlations inferred through LDSC among the cingulo-opercular network (10 volumes).

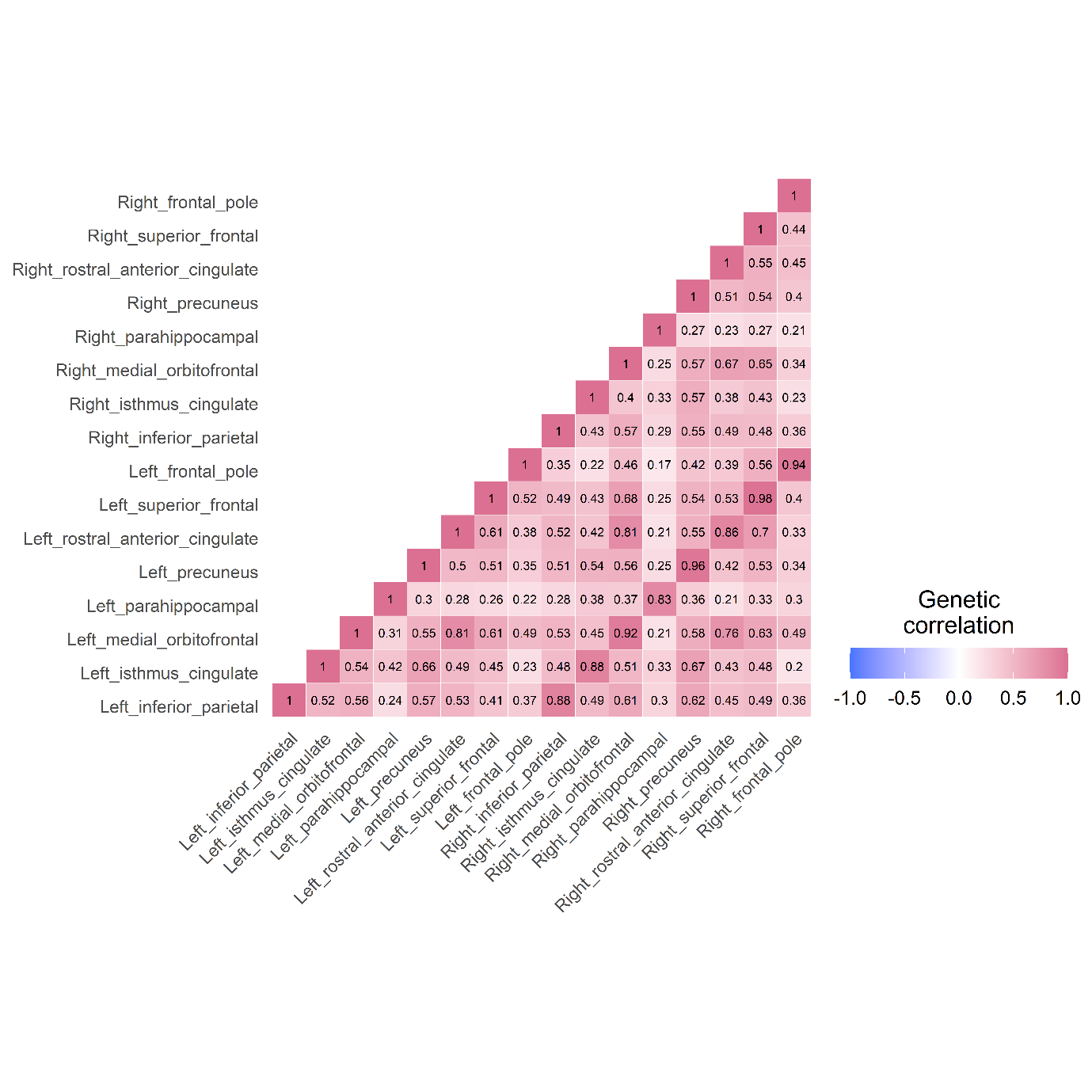

*Supplementary Figure 4.* Genetic correlations inferred through LDSC among the default mode network (16 volumes).

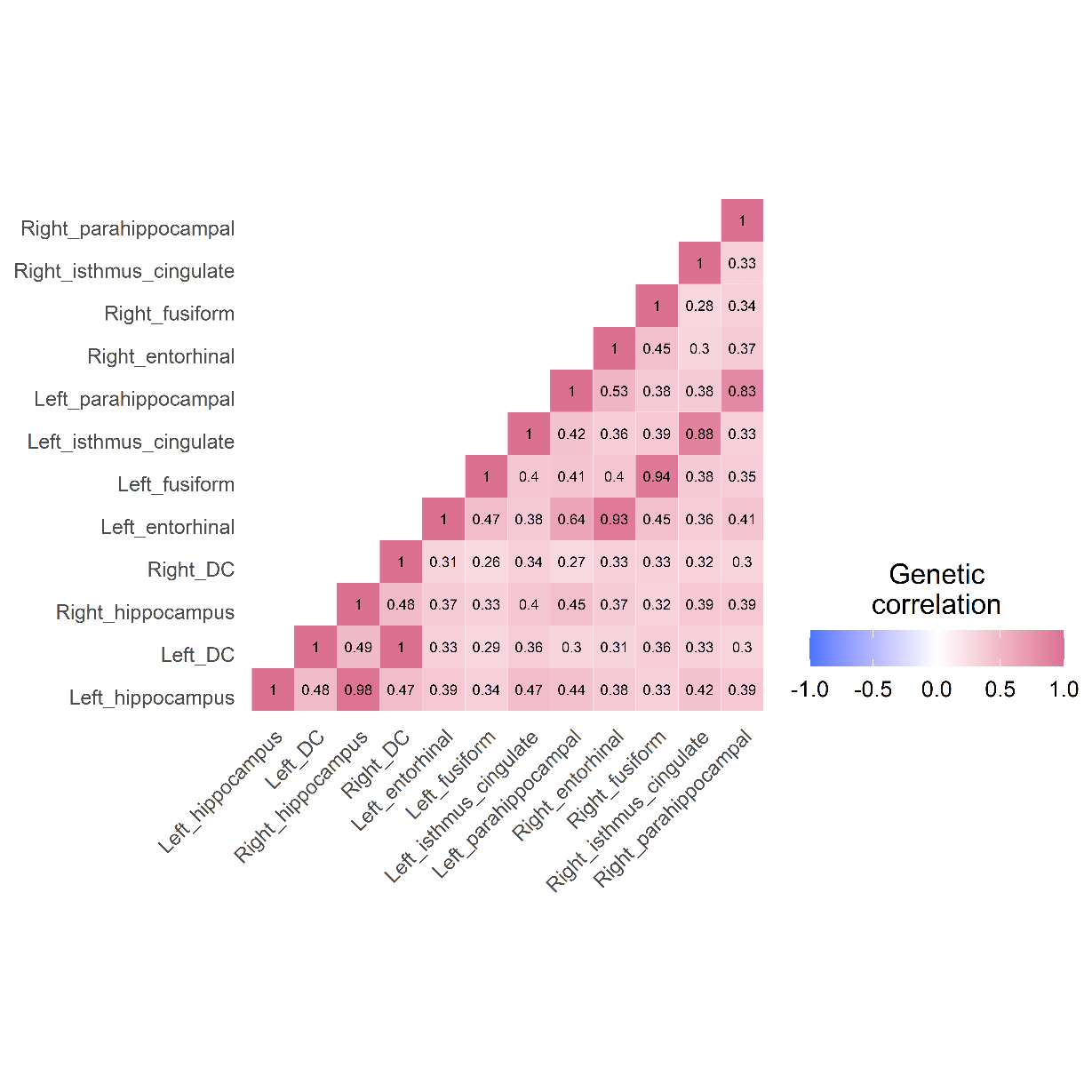

*Supplementary Figure 5.* Genetic correlations inferred through LDSC among the hippocampal-diencephalic network (12 volumes).

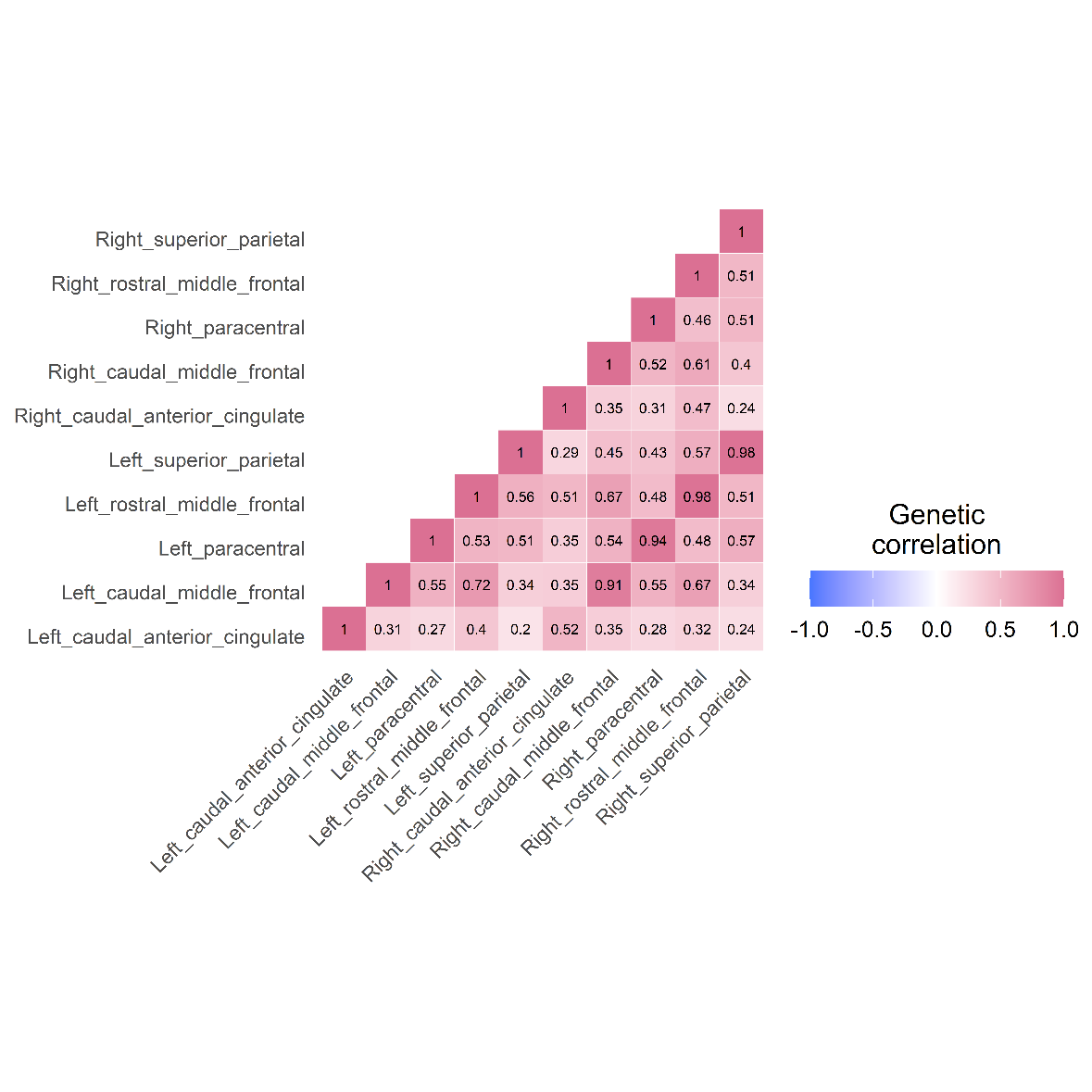

*Supplementary Figure 6.* Genetic correlations inferred through LDSC among the multiple demand network (12 volumes).

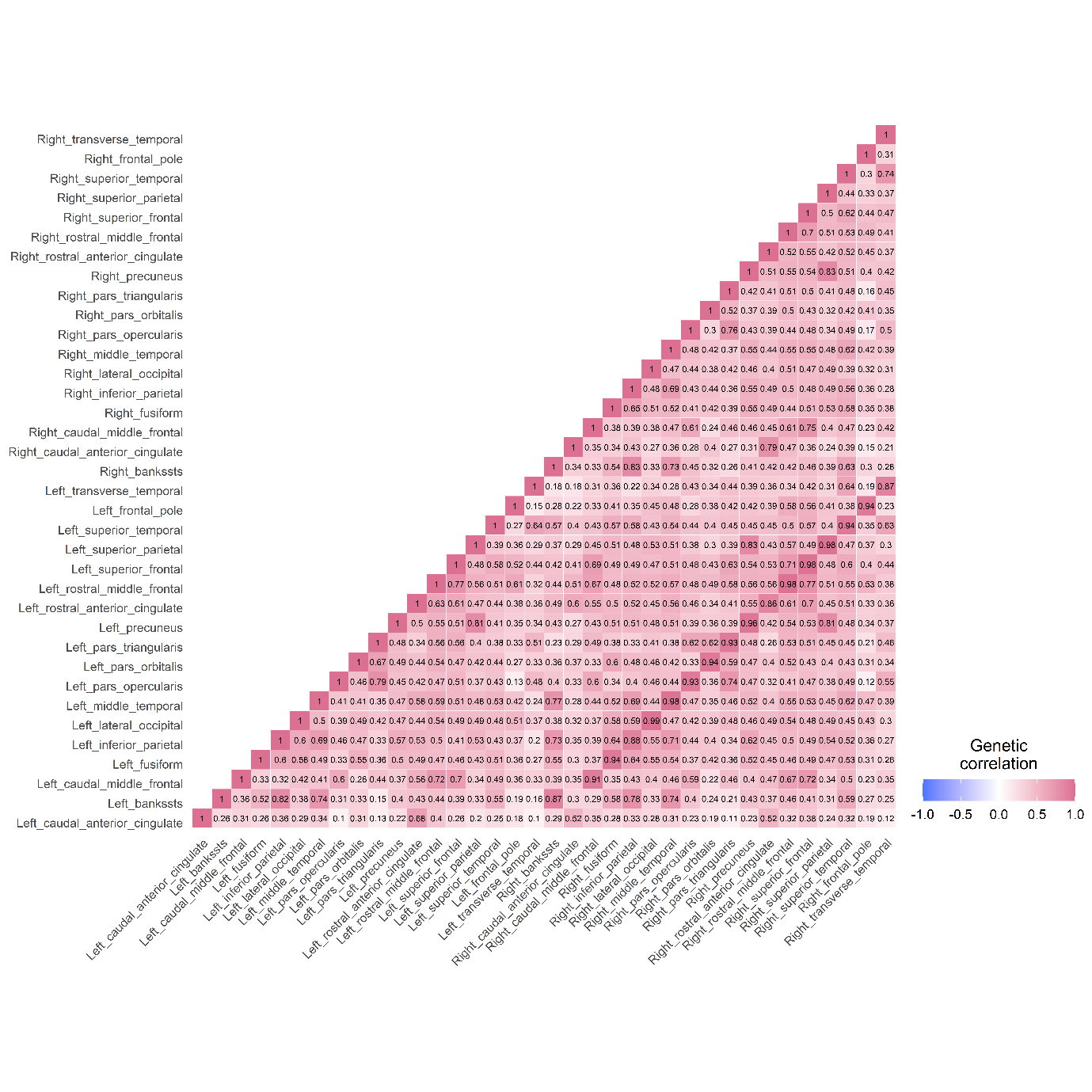

*Supplementary Figure 7.* Genetic correlations inferred through LDSC among the P-FIT network (36 volumes).

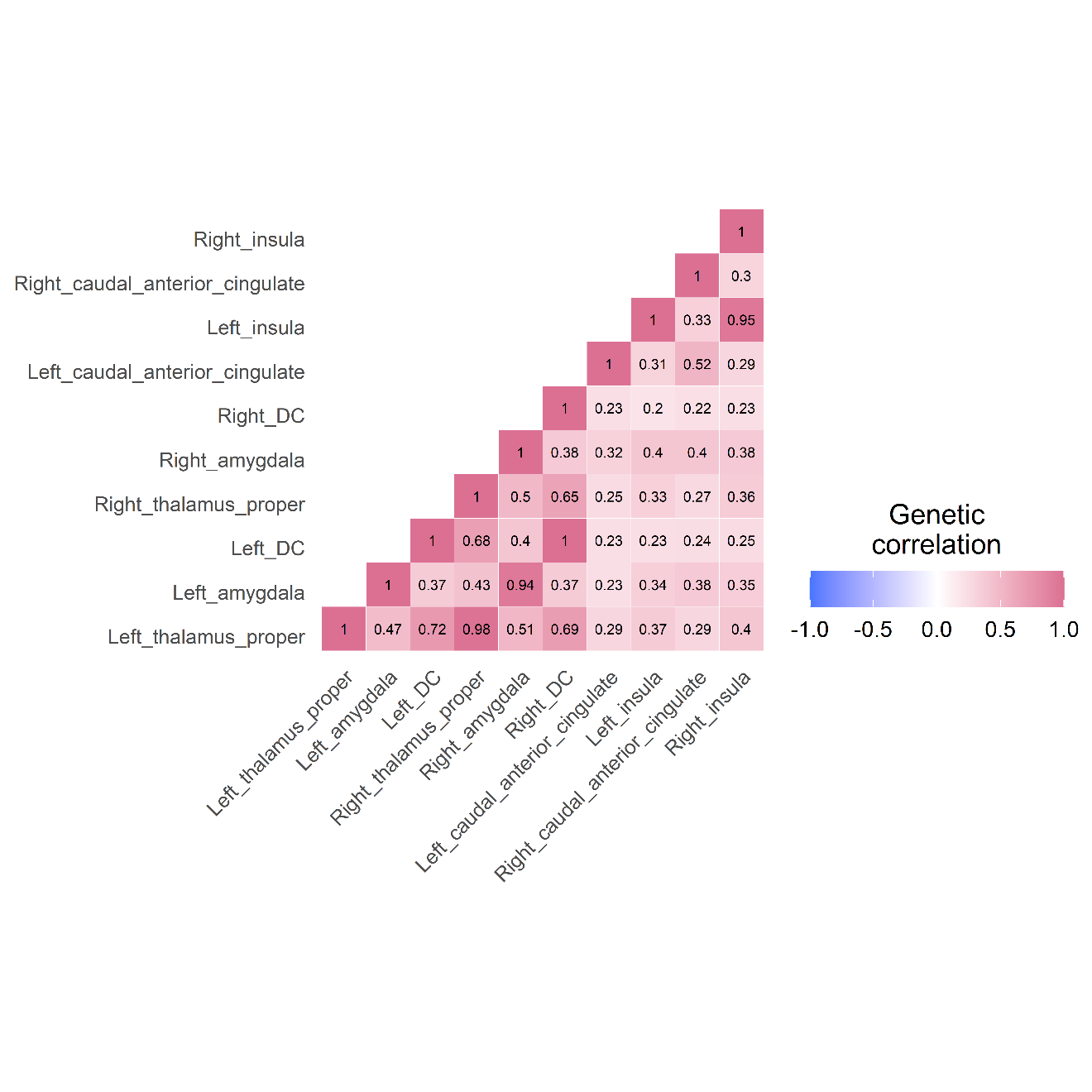

*Supplementary Figure 8.* Genetic correlations inferred through LDSC among the salience network (10 volumes).

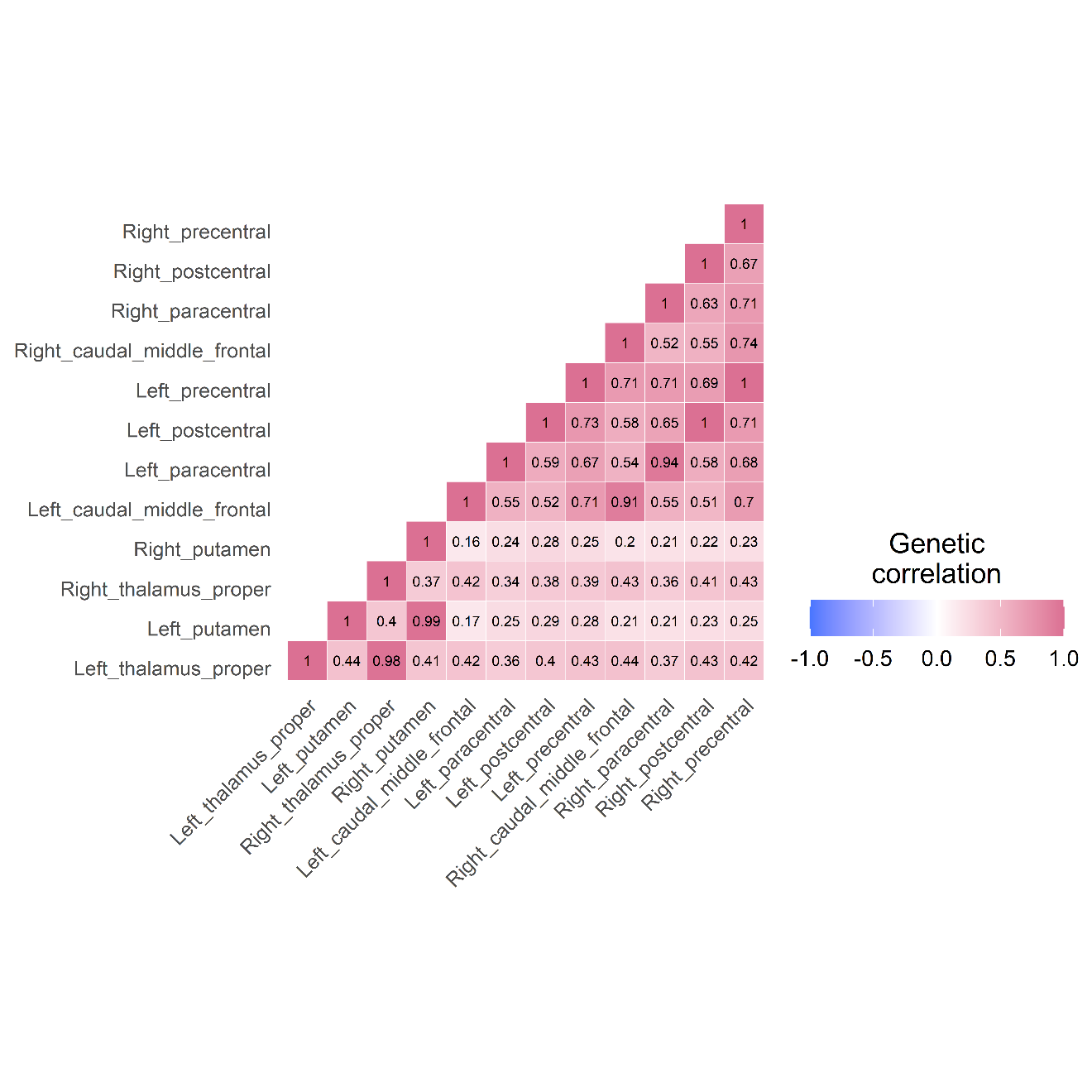

*Supplementary Figure 9.* Genetic correlations inferred through LDSC among the sensorimotor network (12 volumes).

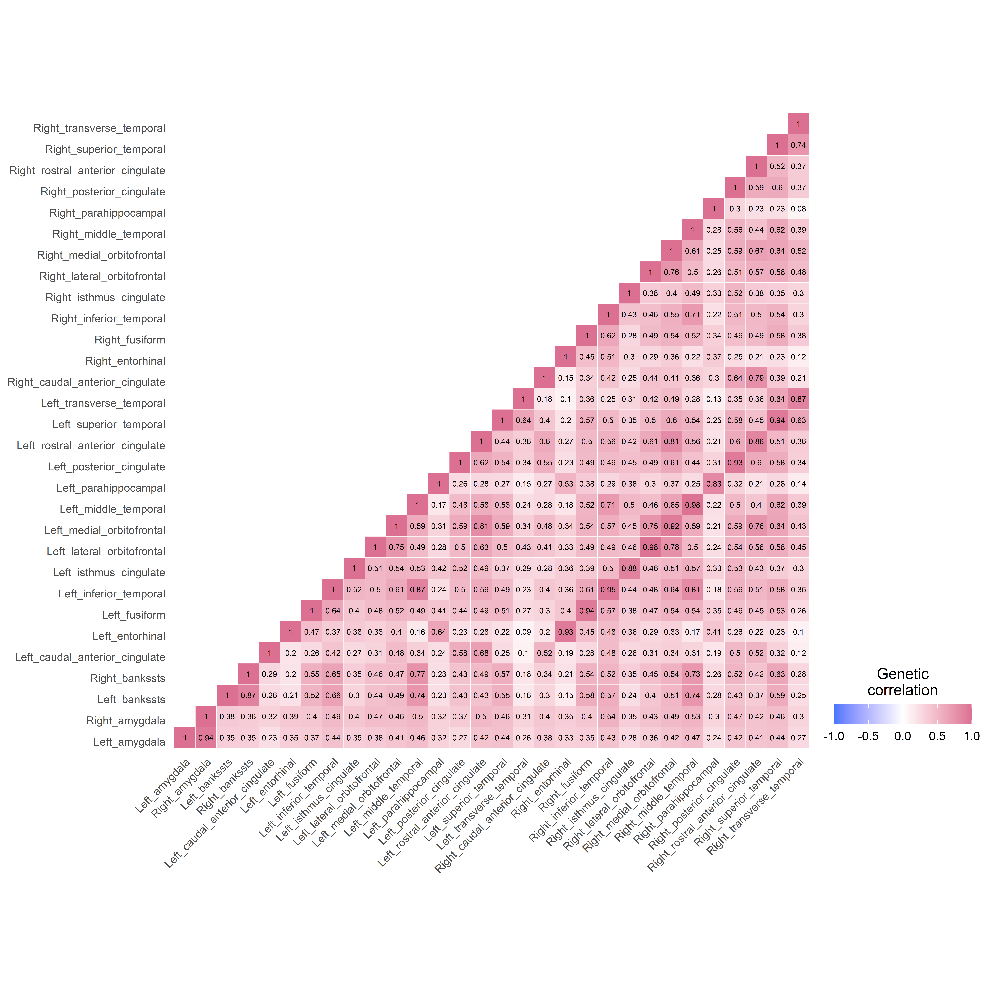
*Supplementary Figure 10.* Genetic correlations inferred through LDSC among the temporo-amygdala-orbitofrontal network (30 volumes).

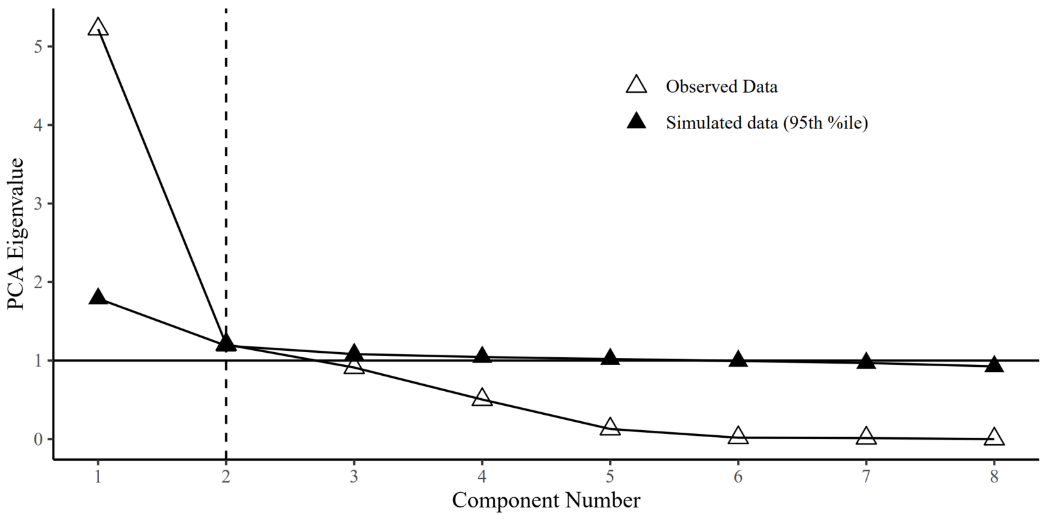
 *Supplementary Figure 11.* Parallel analysis in the central executive network

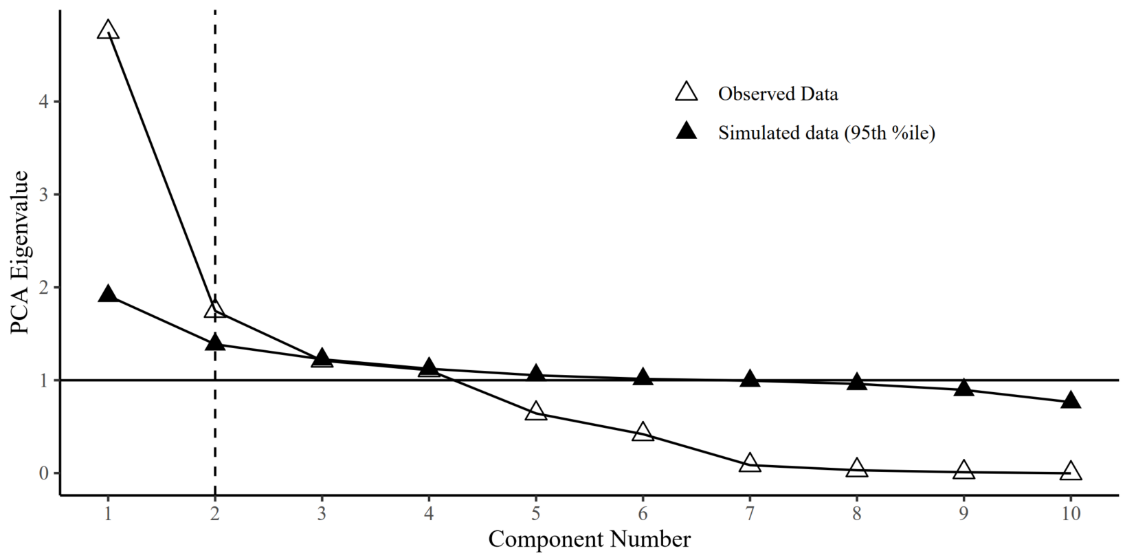
 *Supplementary Figure 12.* Parallel analysis in the cingulo-operular network

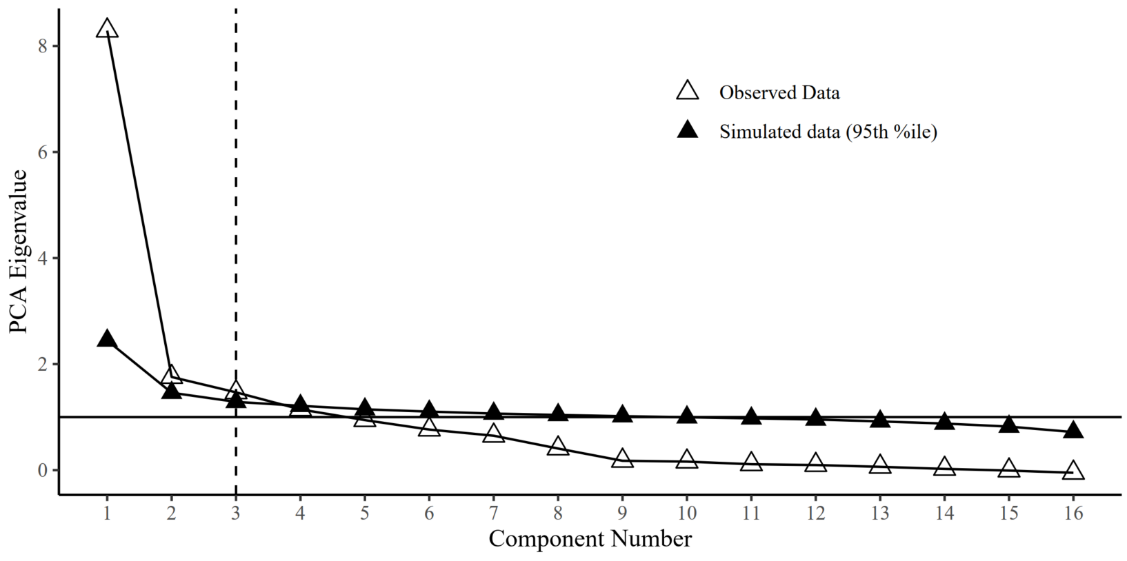
 *Supplementary Figure 13.* Parallel analysis in the default mode network

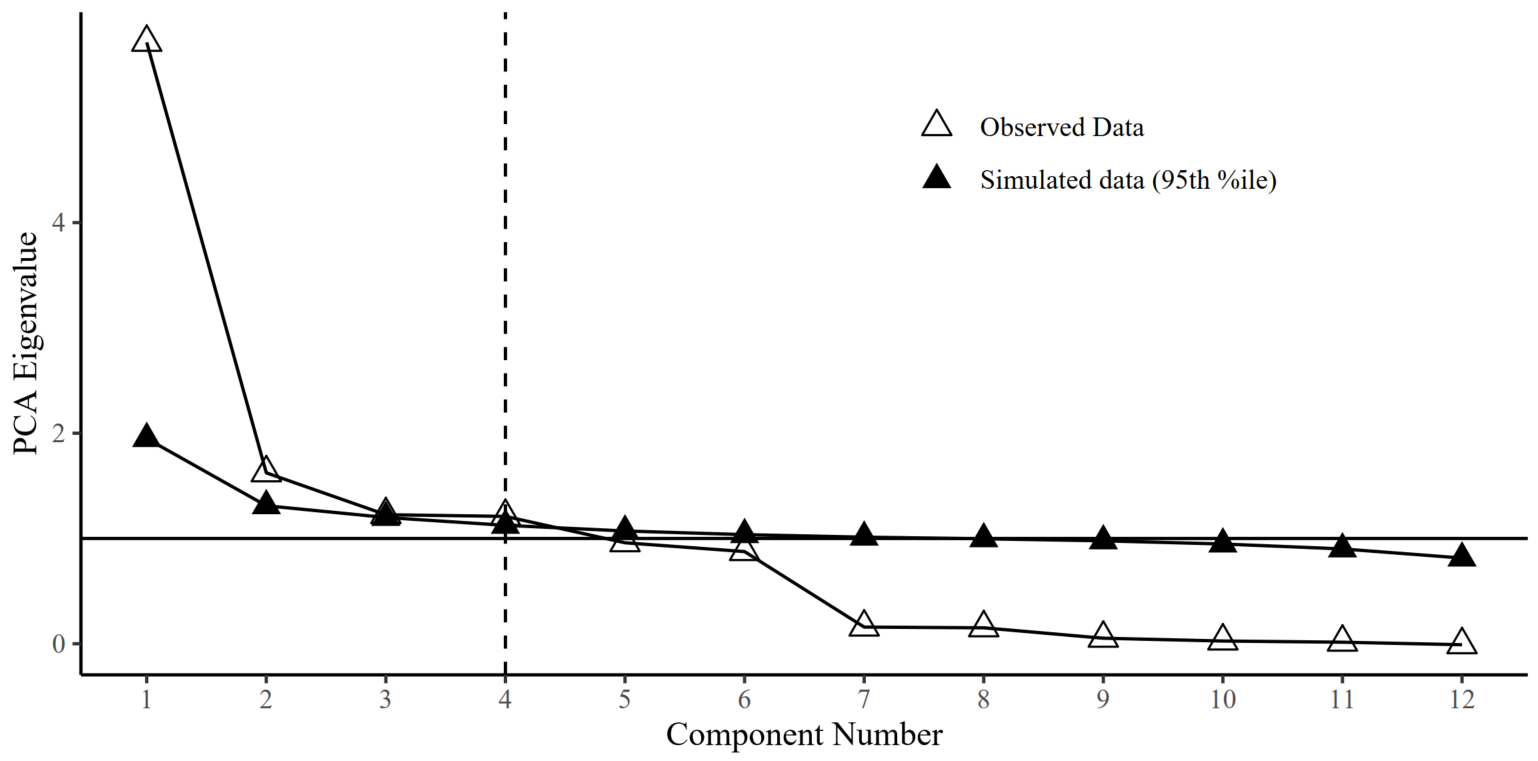
 *Supplementary Figure 14.* Parallel analysis in the hippocampal-diencephalic network

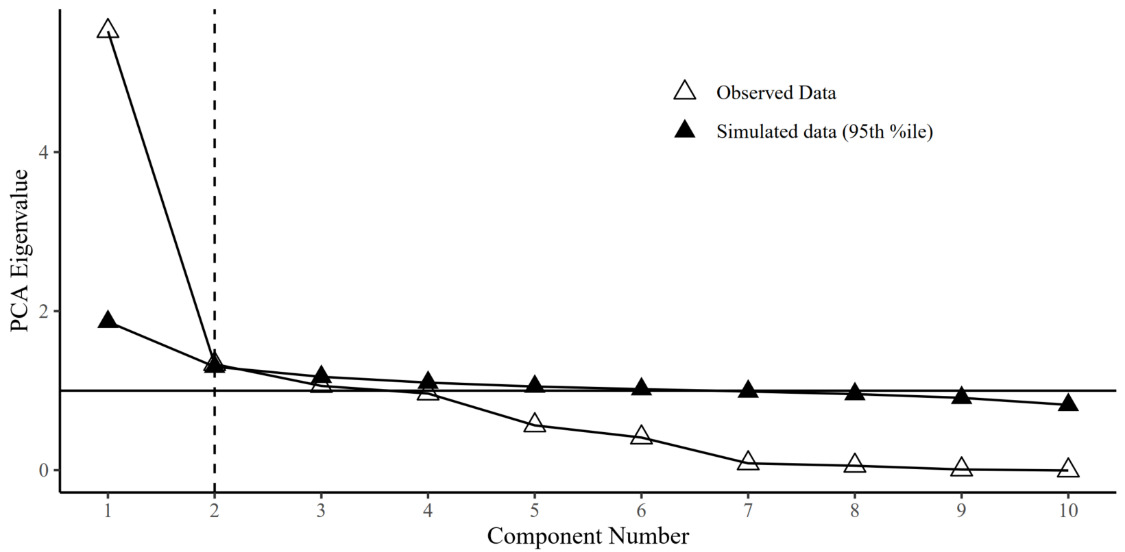
 *Supplementary Figure 15.* Parallel analysis in the multiple demand network

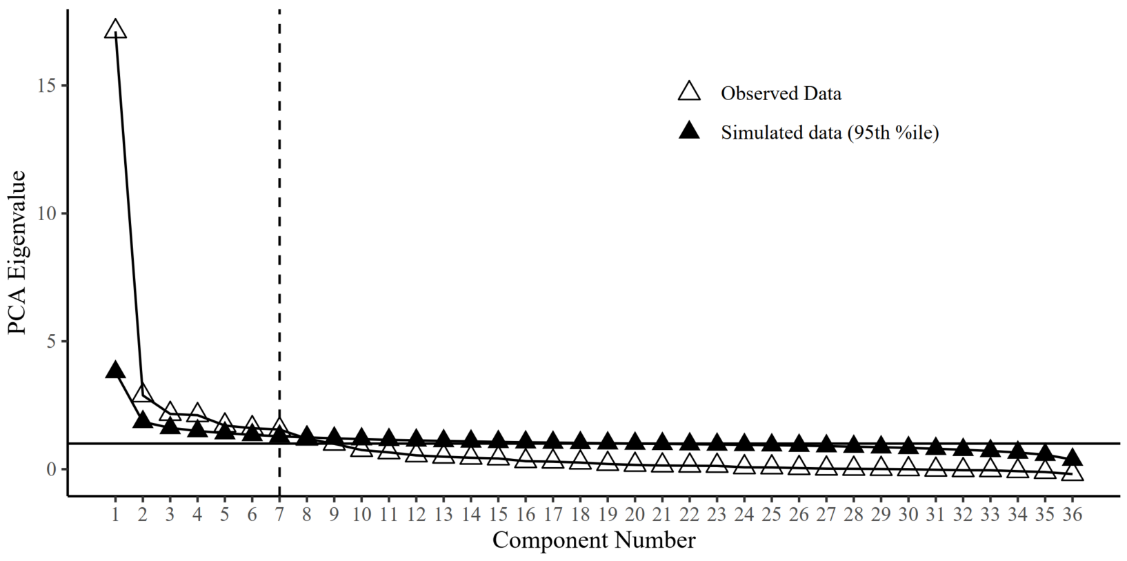
 *Supplementary Figure 16.* Parallel analysis in the P-FIT network

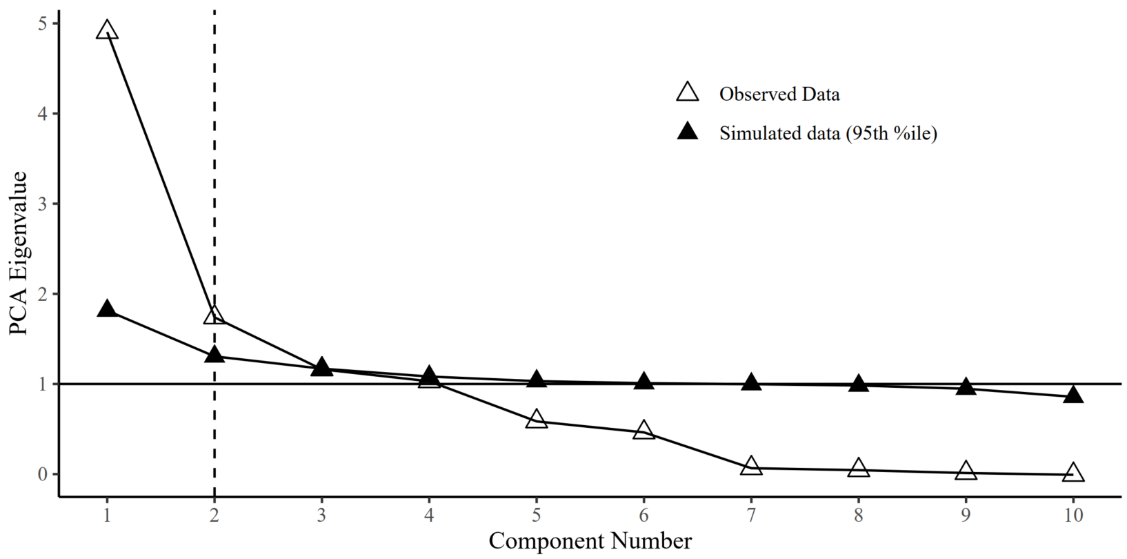
 *Supplementary Figure 17.* Parallel analysis in the salience network

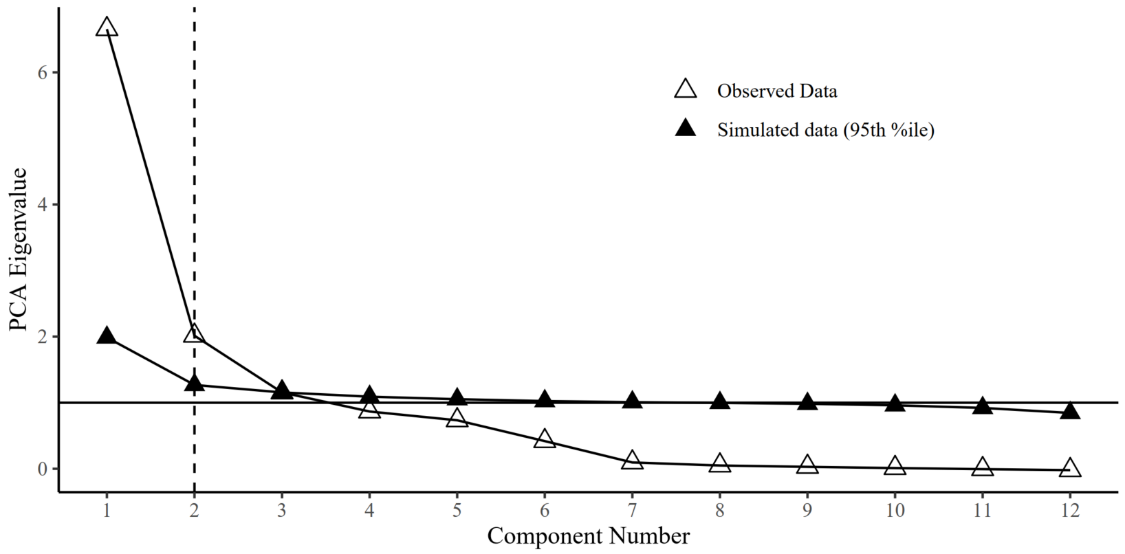
 *Supplementary Figure 18.* Parallel analysis in the sensorimotor network

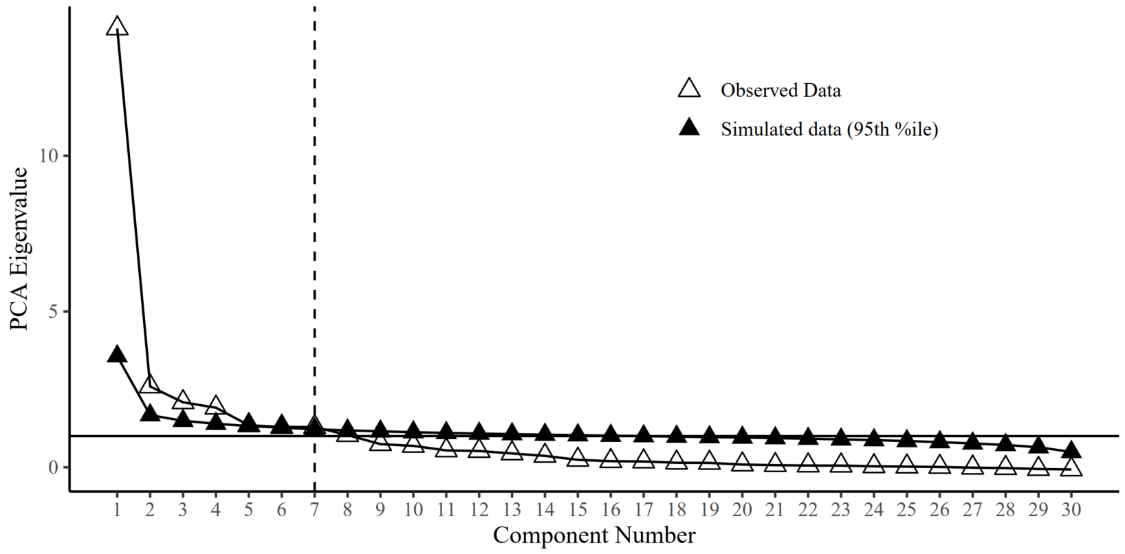
 *Supplementary Figure 19.* Parallel analysis in the temporo-amygdala-orbitofrontal network

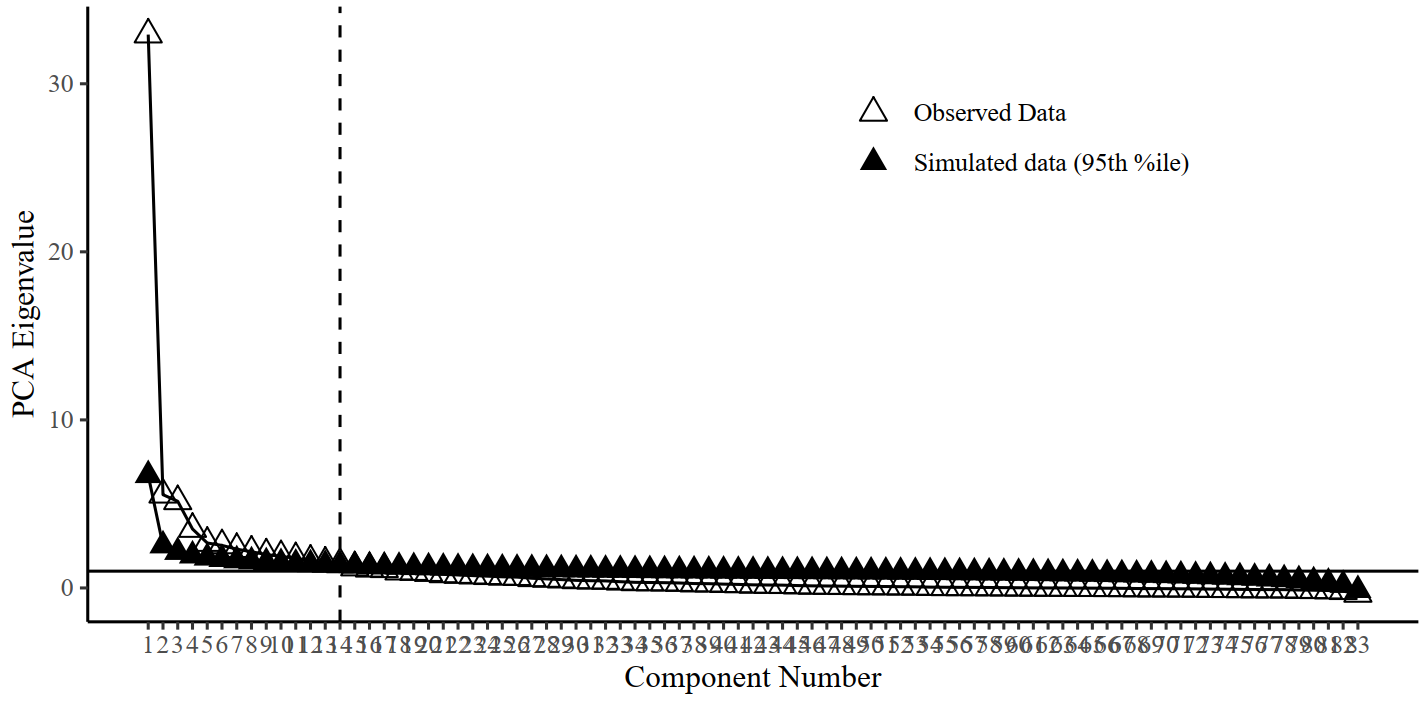

1	14	83

Component Number

*Supplementary Figure 20.* Parallel analysis in the whole brain with 83 nodes.

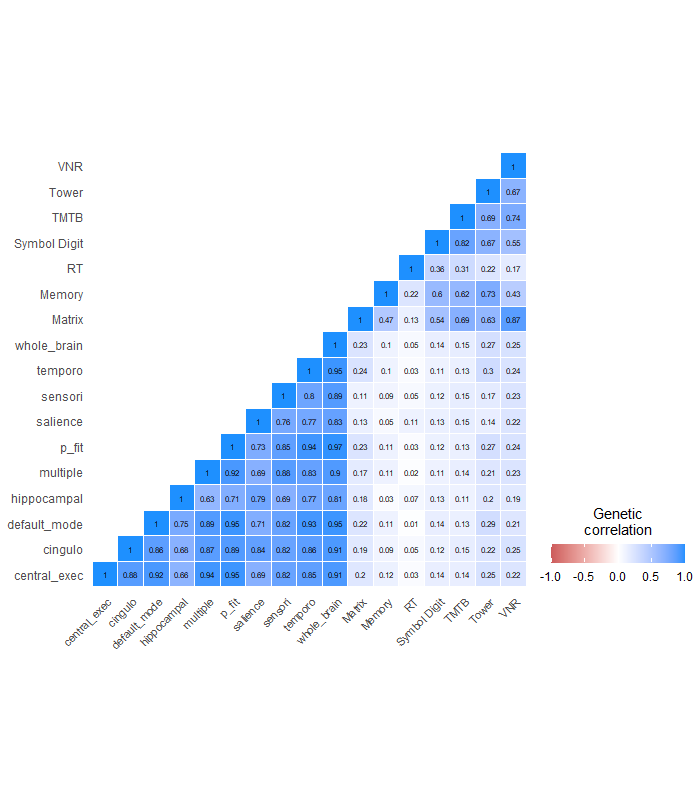

*Supplementary Figure 21.* Genetic correlations between seven cognitive traits and brain networks. Descriptively, performance in the Tower Rearranging Task has the largest association with brain networks in comparison with other cognitive tasks. Abbreviations: Matrix = Matrix Pattern Completion task; Memory = Memory – Pairs Matching Test; RT = Reaction Time; Symbol Digit = Symbol Digit Substitution Task; Trails-B = Trail Making Test – B; Tower = Tower Rearranging Task; VNR = Verbal Numerical Reasoning Test; central exec = central executive; cingulo = cingulo-opercular; hippocampal = hippocampal-diencephalic; multiple = multiple demand; p fit = parieto-frontal integration theory; sensori = sensorimotor; temporo = temporo-amygdala-orbitofrontal

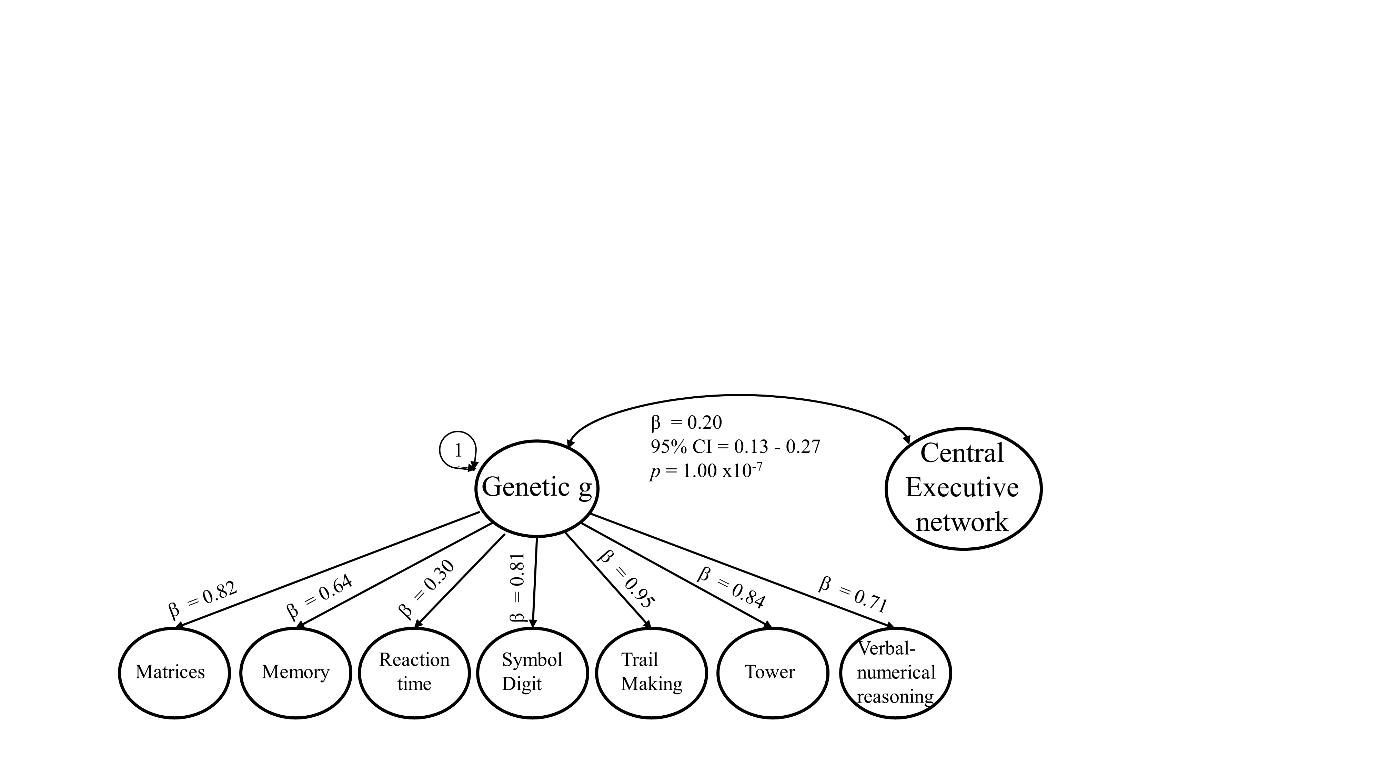

*Supplementary Figure 22.* Genetic correlation between the central executive network and factor *g* modelled for correlation structure of seven cognitive traits. The seven cognitive traits and the network are inferred through LDSC, and the factor through factor analysis. Matrix = Matrix Pattern Completion task; Memory = Memory – Pairs Matching Test; RT = Reaction Time; Symbol Digit = Symbol Digit Substitution Task; Trails-B = Trail Making Test – B; Tower = Tower Rearranging Task; VNR = Verbal Numerical Reasoning Test. Model fit: *χ^2^* = 124.04, df = 20, *p*-value = 2.1 x10^-20^, AIC = 174.04, CFI = 0.97, SRMR = 0.079

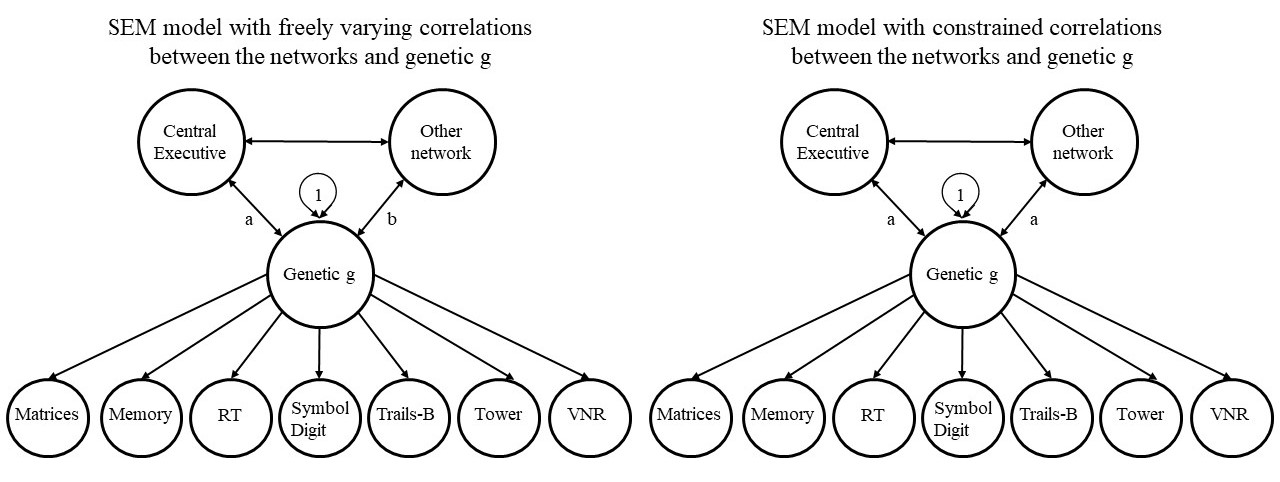

*Supplementary Figure 23.* Illustration of the genomic structural equation models used to test whether correlation magnitudes with genetic general cognitive ability differ between the central executive network and other significantly associated networks. The model on the right freely estimates correlation parameters between two networks and genetic g while allowing for correlations between the networks. In the left model, we force the correlation magnitudes to be the same, and assess whether model fit deteriorates significantly, to conclude whether correlation magnitudes between networks are likely different from each other. The difference between these two models is in the paths leading from the networks to genetic g. In the left model, the paths can take a different value which is why they are labelled with a and b. In the right model, the two paths are specified to be the same and are therefore both labelled with a.

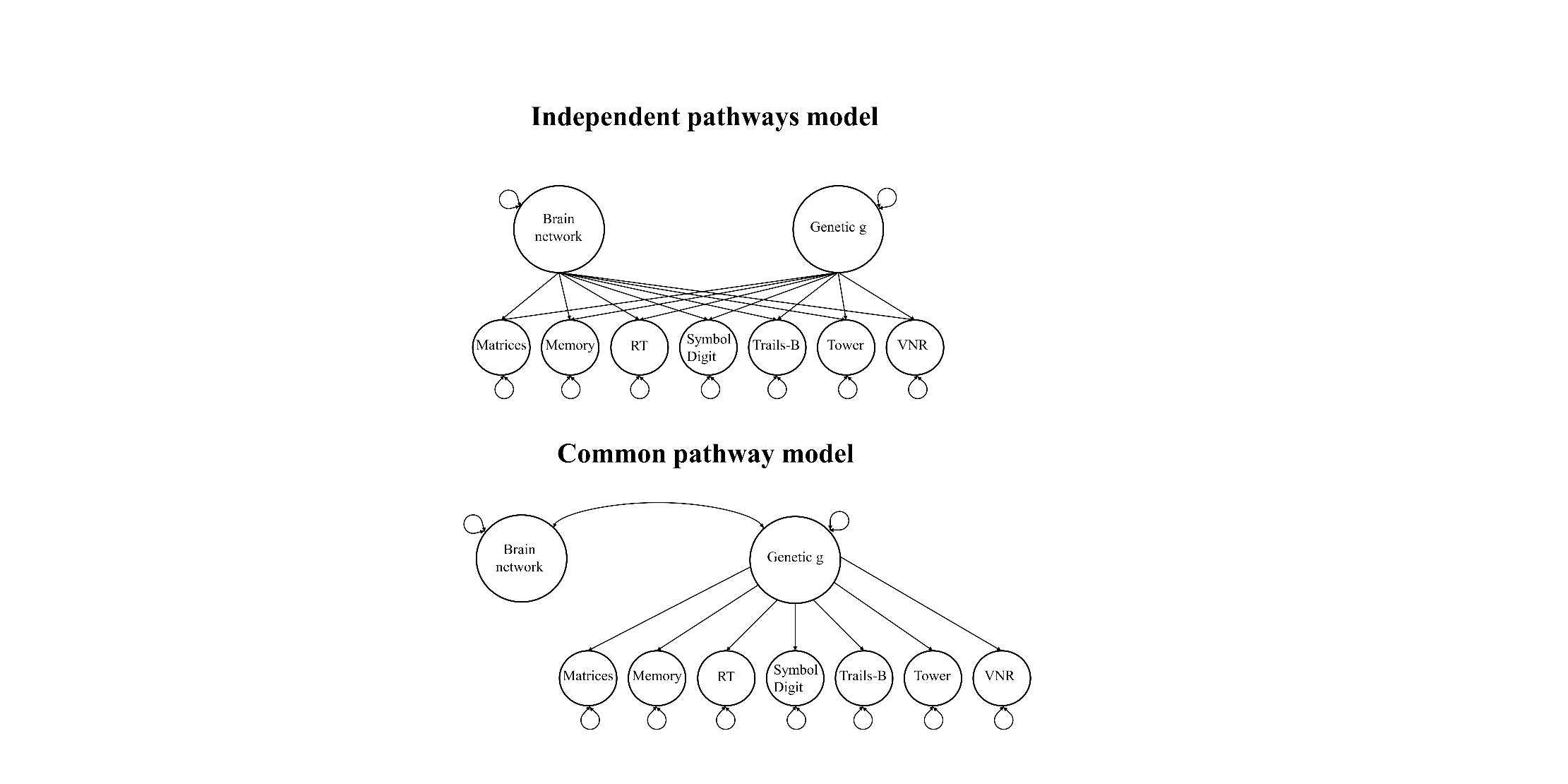

Supplementary Figure 24. Structural equation models to calculate *Q_trait_* heterogeneity indices

### **Supplementary Note**

#### **Further discussion of study limitations**

All limitations that apply to LDSC methodology are relevant to this study. It is known that LDSC estimates can be downwardly biased [55] with larger standard errors as compared with genome-wide complex trait analysis, for example [56]. However, there is no available software allowing genomic structural equation modelling using any of these other methods.

LDSC intercepts should capture some shared phenotypic variation, which could confound the genetic correlation estimates. We found a perfect correlation between intercepts and volume-by-volume phenotypic correlations (*b* = 0.98, SE = 0.009, *p*-value ˂ 2 x10^-16^, *R^2^* = 0.997), indicating that LDSC produced larger intercepts for highly phenotypically correlated traits. This suggests that genetic correlations found in this study are reliable and probably do not rely on methodological artefacts.

LDSC regresses effect sizes from GWAS summary statistics on LD Scores which separates the polygenic signal into signal that correlates with LD (slope), and signal that does not correlate with LD (intercept). Variation captured with the intercept characterises confounding on the cross-trait associations such as sample overlap, population stratification and environmental influences which should be uncorrelated with LD [55, 57]. The correlation between phenotypic associations and intercepts is plausible as it indicates that the intercepts successfully captured and removed confounding effects from the estimate of genetic correlations (slope). We probably obtained genome-wide representative genetic correlations in this study because we discovered pronounced bilateral symmetry of genetic influences, that is near perfect correlations between areas and their homologous counterpart in the opposite hemisphere. This bilateral symmetry was previously reported in twin studies [58, 59] that are sensitive towards overall additive genetic effects and can help inform whether genetic correlations are typical for the whole allelic spectrum.

Another limitation is that precision of genetic correlations depends on the polygenic signal contained in GWAS summary statistics. Low precision and limited systematic variance were linked with unstable genetic correlations [55]. SNP-heritability can be used as a signal-to-noise ratio, to inform whether sufficient polygenic signal was present in a GWAS, and a previous study suggested to exclude traits with SNP-heritability estimates below 0.05 [2]. SNP-heritability estimates of our brain-volumetric traits ranged between 7.6% (SE = 0.01) and 42% (SE = 0.037; median *h^2^* across all volumes = 0.23). The left and right frontal poles had the lowest SNP-heritability estimates, probably because the boundaries of this region are ambiguous [60] resulting in limited systematic variation.
